# Population Imaging of Central Sensitization

**DOI:** 10.1101/2021.11.21.469166

**Authors:** Charles Warwick, Joseph Salsovic, Junichi Hachisuka, Kelly M. Smith, Haichao Chen, James Ibinson, H. Richard Koerber, Sarah E. Ross

## Abstract

Capsaicin applied locally to the skin causes central sensitization that results in allodynia, a state in which pain is elicited by innocuous stimuli. Here, we used two-photon calcium imaging of neurons in the dorsal spinal cord to visualize central sensitization across excitatory interneurons and spinal projection neurons. To distinguish among excitatory neuron subtypes, we developed CICADA, a cell profiling approach that leverages the expression of distinct Gq-coupled receptors. We then identified capsaicin-responsive and capsaicin-sensitized neuronal populations. Capsaicin-sensitized neurons showed emergent responses to low threshold input and increased receptive field sizes consistent with the psychophysical phenomenon that allodynia is observed across an extended secondary zone. Finally, we identified spinal projection neurons that showed a shift in tuning toward low threshold input. These experiments provide a population-level view of central sensitization and a framework with which to model somatosensory integration in the dorsal horn.

**Highlight:** Warwick et al. use two-photon calcium imaging coupled with pharmacological profiling to identify neuronal populations in the spinal dorsal horn that mediate capsaicin-induced central sensitization.

## Introduction

Persistent noxious input drives changes in central circuitry that alters the integration of somatosensory stimuli, resulting in abnormally amplified pain [1]. This central sensitization can be protective in the context of acute injury because it engenders hypervigilance during recovery. In the context of chronic pain, however, central sensitization is maladaptive, leading to suffering. Given the link between central sensitization and the chronification of pain, it is important to understand activity-induced changes in the central processing of nociception.

Capsaicin-induced hypersensitivity is a model of central sensitization that is well characterized in psychophysical [2, 3] and electrophysiological studies [4, 5]. In this model, cutaneous application of capsaicin activates TRPV1^+^ C-fibers resulting in burning pain that lasts minutes, and central sensitization in the dorsal horn that lasts hours. Two salient features of this central sensitization are that low threshold mechanical stimuli, which are normally innocuous, are perceived as painful (allodynia), and that the area of allodynia (the secondary zone) is much larger than the area of capsaicin-treated skin. In light of these findings, capsaicin-induced allodynia is thought to be a central phenomenon, likely due to plasticity at the level of the spinal cord [5–7]. Consistent with this idea, electrophysiological studies have revealed there are synaptic changes in spinal projection neurons after capsaicin treatment [4, 8]. However, the identity of which spinal populations are involved in generating this plasticity have yet to be examined.

Ca^2+^ imaging is a powerful approach to visualize neural activity across a network. However, there are very few studies to date that have applied this method to the spinal cord [9, 10] in part because of difficulty in mitigating respiratory movements and a high degree of neuronal diversity [11]. To overcome these limitations, we used an *ex vivo* approach to facilitate the imaging and developed a novel strategy of cell type identification to empower the interpretation. In the *ex vivo* somatosensory preparation, the skin, nerves, dorsal root ganglia, and spinal cord are dissected out in continuum, allowing the application of quantitative sensory stimuli to the skin. This reduced preparation is advantageous because it provides superior image quality and stability, eliminates the confound of anesthesia, and facilitates extensive pharmacological manipulation. Additionally, rather than imaging a mixed population of unidentified neurons, we focused on excitatory neurons, and further distinguished interneurons from projection neurons by virtue of retrograde labeling from the lateral parabrachial nucleus. To further differentiate the excitatory populations, we leveraged the heterogeneous expression Gq-coupled G-protein coupled receptors (GPCRs). Because activation of Gq-coupled GPCRs gives rise to the release of Ca^2+^ from internal stores, the application of GPCR ligands allows us to visualize which cells express which receptors. We have called this approach CICADA for **C**ell-type **I**dentification by **C**a2+-coupled **A**ctivity through **D**rug **A**ctivation.

Here, we used 2-photon Ca^2+^ imaging in the superficial dorsal horn to define the response properties of both spinoparabrachial (SPB) neurons and excitatory spinal interneurons to quantitative mechanical and thermal stimuli. Using CICADA, we defined seven excitatory interneuron populations based on their responses to Gq-coupled GPCR agonists. We then determined how central sensitization alters their response properties and identified distinct CICADA-defined populations that comprise capsaicin-responders and capsaicin-sensitized populations. Within identified capsaicin-sensitized populations, some show selectively amplified responses to low threshold input suggestive of allodynia, whereas others show increases in receptive field size consistent with a secondary zone of allodynia expanding beyond the capsaicin injection site. Finally, we show that capsaicin-induced central sensitization alters the response properties of SPB neurons, which show a shift in tuning towards low threshold input. Together, these data represent a population view of central sensitization in the superficial dorsal horn.

## Results

### *Ex vivo* Ca^2+^ imaging in the spinal dorsal horn

We performed two-photon Ca^2+^ imaging of excitatory neurons in the superficial dorsal horn of the spinal cord in response to natural cutaneous stimulation (Fig. 1a). For these recordings, we used an *ex vivo* somatosensory preparation in which the spinal cord, nerves (saphenous and lateral femoral), and skin (dorsal hind paw and proximal hip) were dissected in continuum (Fig. 1b). This approach allowed us to manipulate sensory input at the skin with mechanical, thermal, and chemical stimuli. Excitatory neurons were selectively labeled using the *Vglut2-ires-Cre* allele together with Ai96, enabling Cre-dependent expression of GCaMP6s from the Rosa locus. SPB neurons were identified by back-labeling from the lateral parabrachial nucleus with a vital, red-fluorescent dye, DiI (Fig. 1c). To record activity across neurons of different laminae, we imaged from multiple optical planes simultaneously (Fig. 1d). In this study, we focused on neurons ranging from 0 – 60 µm below the dorsal surface of the spinal cord (lamina I, IIo and IIi), but this approach allows us to visualize deeper laminae, extending down to at least 150 µm from the surface of the spinal cord (Extended Data Fig. 1). Within a 528 x 221 µm field of view, this labeling and imaging strategy enabled the visualization of ∼300 excitatory neurons and ∼6 to 8 SPB neurons per animal. In total, 11 mice were imaged with 2,914 individual cells recorded across all sets of experiments. Light brushing of the skin gave rise to Ca^2+^ transients in a subset of neurons that was highly reproducible (Fig. 1e,f) and showed no significant rundown of response after repeated stimulation (Extended Data Fig. 2).

**Fig. 1.**
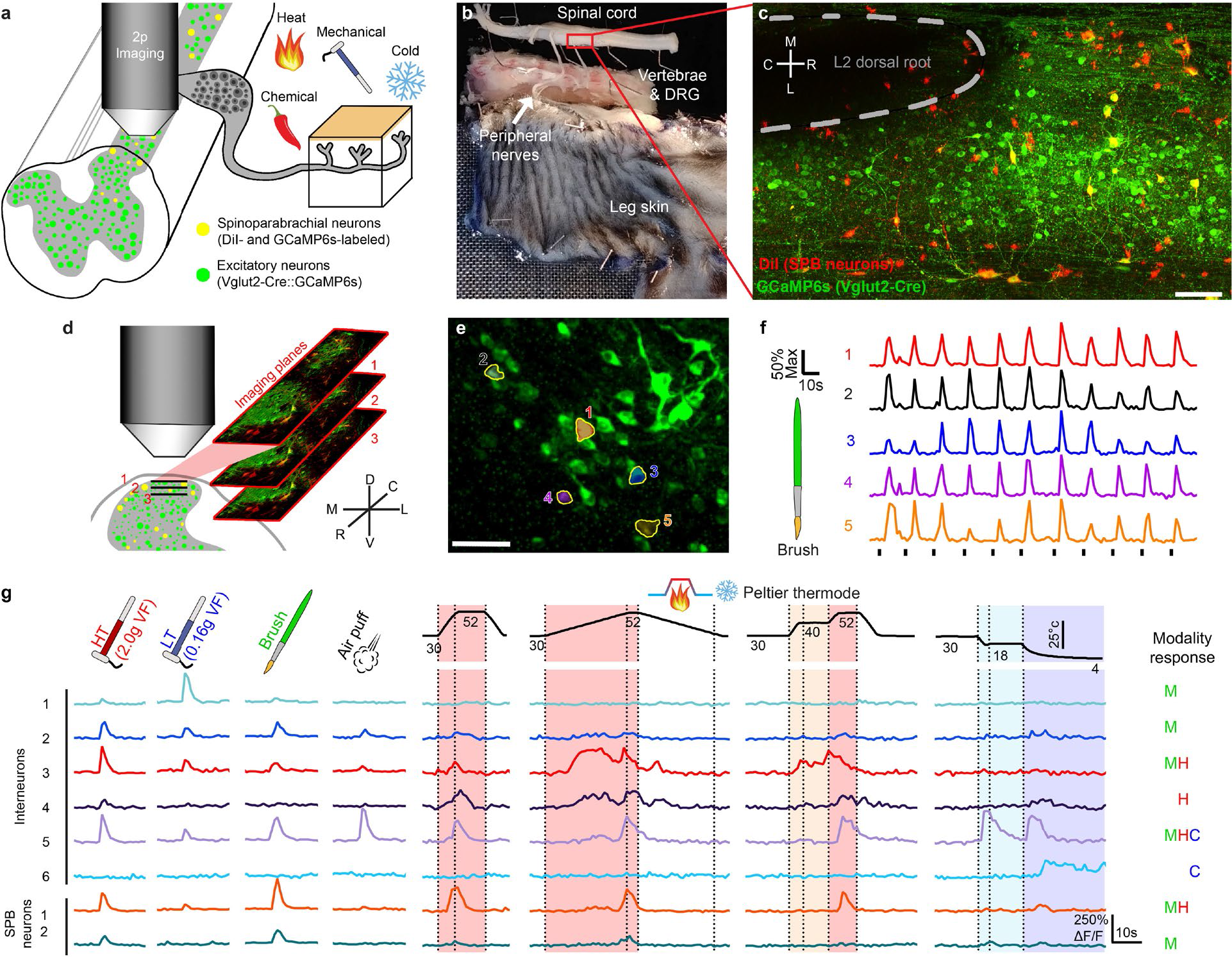
Population imaging in the *ex vivo* somatosensory preparation. **a**, Schematic of the *ex vivo* somatosensory preparation and neuronal labeling strategy. **b**, Picture of the *ex vivo* somatosensory preparation with a typical recording region rostral of the L1-L2 dorsal root entry zone highlighted in red. **c**, Low-magnification fluorescence image of the L2 dorsal root entry zone (scale bar, 100 µm) showing GCaMP6s expression in *Vglut2-Cre* positive excitatory neurons (green) and SPB neurons (red). **d**, Schematic of the multi-plane (XYZT) imaging protocol used for Ca^2+^ imaging. 3 separate optical planes with a spacing of 14 µm are imaged simultaneously allowing parallel access to lamina I and II neurons. **e**, Example maximum ΔF/F image with 5 brush responsive cells circled (scale bar, 50 µm). **f**, ΔF/F Ca^2+^ traces from the 5 cells circled in **e** showing consistent responses to repeated brushing of the skin. **g**, Top: stimulation type for mechanical and thermal stimuli. High threshold (HT) and low threshold (LT) stimuli were applied serially across an area of skin 15×15 mm, while all other stimuli covered the entire area with each trial. Bottom: Representative ΔF/F Ca^2+^ traces from both interneurons and SPB neurons in response to the indicated cutaneous stimuli. Responsiveness to cooling (C) mechanical stimulation (M) or heating (H) is indicated on the right.

To characterize the functional response properties of excitatory neurons in the dorsal horn, we applied a series of mechanical and thermal stimuli to the skin. For static mechanical stimulation, we used von Frey filaments of 2.0 g for high threshold (HT) or 0.16 g for low threshold (LT). For dynamic mechanical stimulation, we applied a wide brush or an air puff (60 PSI through a 0.8 mm nozzle). Thermal stimuli consisted of fast and slow ramps to warm (40°C), hot (52°C), cool (18°C), or cold (4°C) temperatures using a Peltier thermode. Examples of the Ca^2+^ responses in several excitatory interneurons and SPBs are shown (Fig. 1g), illustrating that individual neurons have complex responses to diverse sensory inputs. In general, we found that ∼70% of GCaMP6s-expressing neurons within the field of view responded to one or more of these natural stimuli.

### Polymodality and tuning

As a first step to functionally classify neurons in the dorsal horn, we defined a responder as a cell showing a stimulus-evoked Ca^2+^ transient with a peak amplitude that was at least 125% ΔF/F and more than 6 standard deviations above baseline activity (Fig. 2a and Extended Data Fig. 3). Of cells that responded to cutaneous stimuli, we found that 86% of excitatory interneurons responded to mechanical, 58% to heat, and 28% to cold (Fig. 2b). Among SPB neurons these numbers were even higher: 100% responded to mechanical, 52% to heat, and 47% to cold. Indeed, compared to interneurons, the fraction of SPB neurons that responded to all three cardinal modalities (mechanical, heat, and cold) was significantly higher, indicating a greater degree of modality convergence (Fig. 2c). Across mechanical sub-modalities, we observed that HT (2.0 g), LT (0.16 g), and light brushing stimuli activated 82%, 59% and 37% of superficial interneurons, and 100%, 52% and 53% of SPB neurons, respectively (Fig. 2d). Again, the proportion of SPB neurons that responded to all three mechanical stimuli (pan-mechanical) was significantly higher than that of interneurons (Fig. 2e). Thus, many spinal interneurons and even more SPB neurons show polymodal responses to cardinal sensory modalities and respond broadly to a variety of subtypes of mechanical stimuli.

**Fig. 2.**
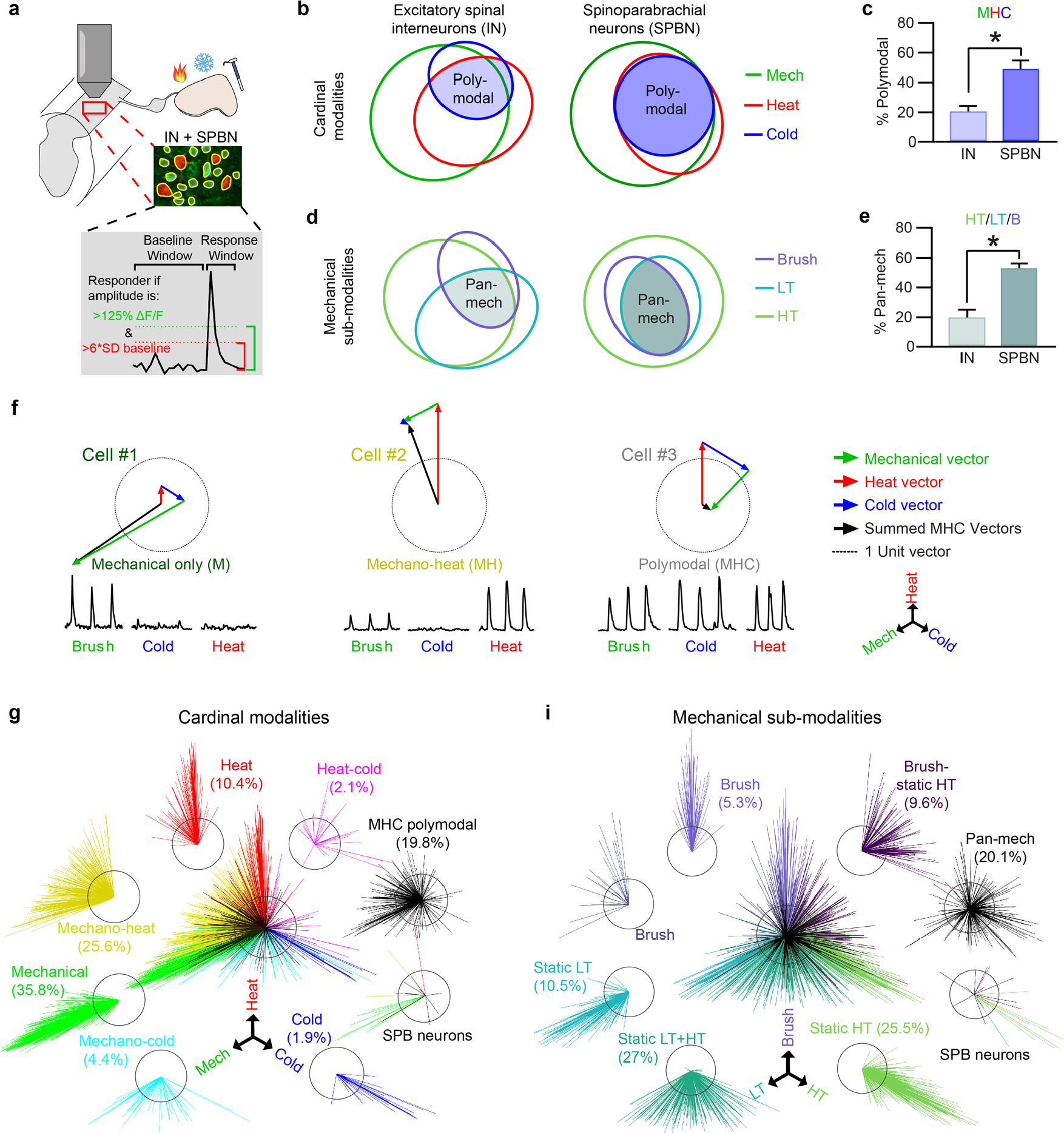
Polymodality and tuning among excitatory interneurons and projection neurons. **a**, Experimental design of imaging (top) and schematic of response thresholds (bottom). **b**, Proportions of excitatory interneurons (IN, left) and SPB neurons (SPBN, right) that were responsive to cardinal modalities (cold, mechanical, heat). **c**, Comparison of cardinal polymodality within INs and SPBNs. There was a significant main effect of modality (*F* (7, 35) = 4.447), *P*=0.0013, and an interaction between modality and cell type (*F* (7, 27) = 2.725), *P*= 0.0283 (RMLM model). Post hoc testing was performed between cell types for each modality, \**Q*<0.05. **d**, Proportions of INs (left) and SBPNs (right) that were responsive to the mechanical sub-modalities: brush (purple), static low threshold (LT, blue), or static high threshold (HT, green). Pan-mechanical sensitivity (pan-mech) indicates responsiveness to all tested types of mechanical stimuli. **e**, Comparison of pan-mechanical prevalence in INs and SPBNs. There was a significant main effect of modality (*F* (6, 18) = 7.372), *P*=0.0004, and an interaction between modality and cell type (*F* (6, 18) = 4.582), *P*= 0.0054 (2-way RM ANOVA). Post hoc testing was performed between cell types for each modality, \**Q*<0.05. **f**, Schematic and examples of vector-based tuning. Top: ΔF/F Ca^2+^ traces evoked by brush (green), cold (blue), and heat (red). Bottom: modality response vectors (colored) and summed vectors indicating the final tuning (black). **g**,**h** Cardinal (**g**) and mechanical sub-modality (**h** ) tuning of SDH interneurons grouped by their threshold-based categorization. SPBNs are shown separately. *N* = 1265 cells for the cardinal modalities and 1080 cells for the mechanical sub-modalities from 5 mice including SPB neurons and 6 mice for interneurons.

Although excitatory neurons were frequently polymodal, we nevertheless observed differences in the magnitude of responses to each modality/sub-modality. To examine this observation in more detail, we developed a two-dimensional vector-based method to visualize and quantify sensory tuning (Fig. 2f). For each cell, three vectors were calculated based on the amplitude of the cell’s responses to mechanical, heat, and cold stimuli, with vectors for each modality plotted at 120° angles from one another. The overall tuning of a cell was determined by summing these three vectors, giving rise to a single line in which the direction indicates the relative modality preference and the length indicates the magnitude of preference. Applying this vector-based tuning, we found that although many cells are polymodal they nonetheless show tuning preferences across cardinal modalities (Fig. 2g) and mechanical sub-modalities (Fig. 2h).

### Somatotopy and functional organization

We know that the cells within the dorsal horn are organized somatotopically based on *in vivo* single unit recordings [12–14], but how this organization is manifest at a population level has never been functionally visualized. To examine this feature, we probed the skin with a LT filament at proximal, intermediate, or distal regions (Fig. 3a). This receptive field mapping showed that the stimulation of proximal skin activated neurons that are more lateral within the dorsal horn, whereas stimulation of a distal skin activated neurons that are more medial (Fig. 3b,c). These data are consistent with the idea that the spinal cord is somatotopically organized such that information pertaining to a given area of skin is mapped across the medio-lateral and rostro-caudal axis with modality-relevant information distributed across the dorsal-ventral axis (Extended Data Fig. 4). Next, we examined how the intensity of stimulation affected receptive field size by stimulating at 16 sites equally spaced in a 4 x 4 grid using a HT, LT, or airpuff (Fig. 3d). We observed that the receptive field size of individual excitatory neurons was significantly affected by the stimulus intensity: the determined receptive field size with a HT filament was significantly greater than that with a LT filament, which was significantly greater than that with airpuff (Fig. 3e-f). We also found that neurons that were closer to the surface (*i.e.*, lamina I) had significantly larger receptive fields than neurons that were deeper (i.e., lamina II) (Fig. 3g-i).

**Fig. 3.**
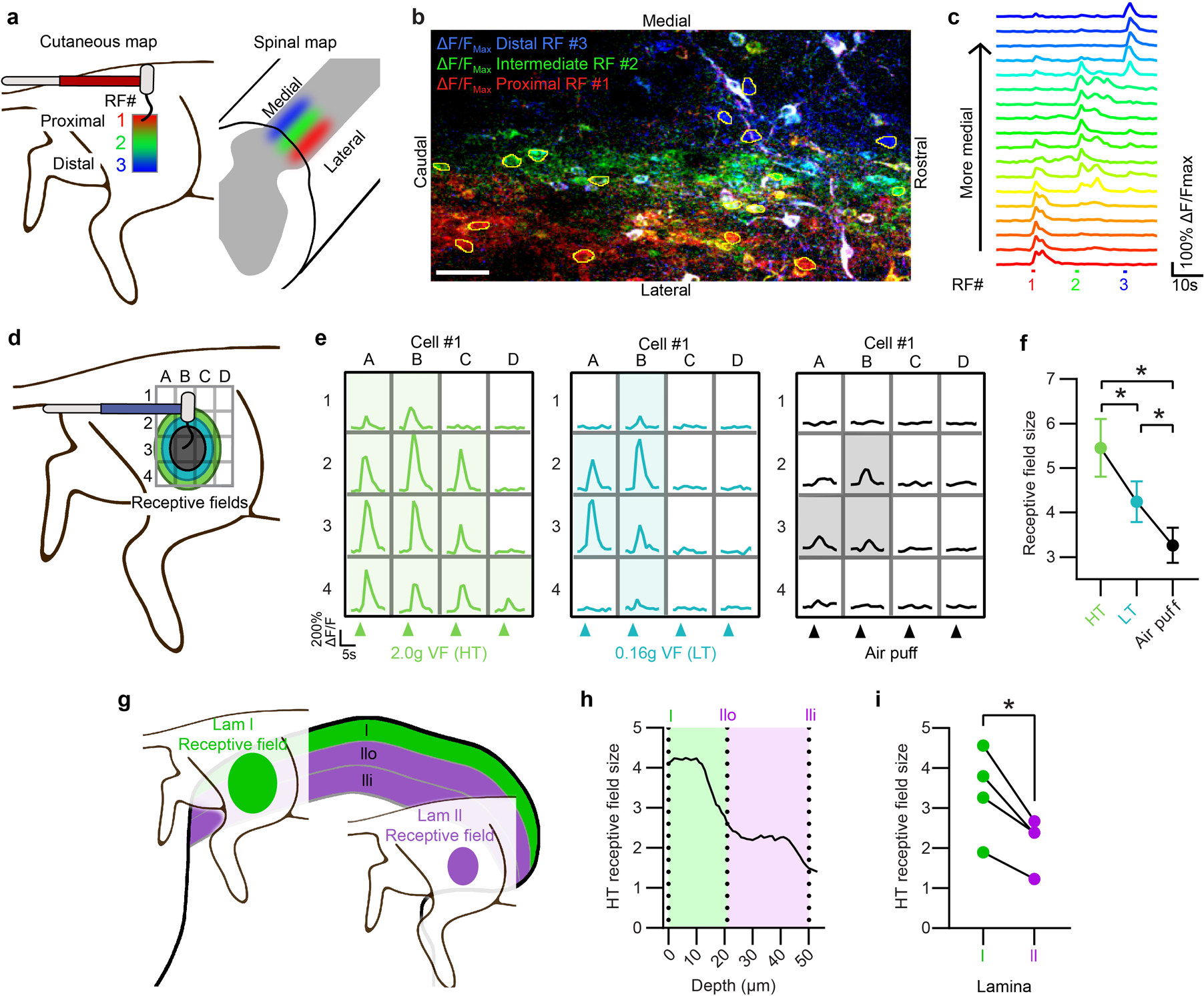
Receptive fields in SDH are shaped by somatotopy, modality, and relative dorso- ventral position. **a**, Schematic illustrating proximal-distal somatotopy. **b**, Field of view in the superficial SDH showing maximum intensity responses after LT stimulation pseudo-colored to each of the indicated receptive fields (RF) in **a**: red, proximal; green, intermediate; blue, distal. **c**, ΔF/F Ca^2+^ traces from circled cells in **b**, showing responses to stimulation at the three indicated RFs, serially stimulated from proximal to distal. Traces are arranged by their somatic medio-lateral location. **d**, Schematic illustrating RF sizes for different mechanical stimuli: HT, green; LT, blue; and air puff, black. **e**, Representative ΔF/F Ca^2+^ traces at 16 RFs for each stimulation type. **f**, Mean RF size of excitatory SDH neurons responsive to all 3 modalities. There was a significant main effect of stimulation type (*F* (1.686, 227.6) = 49.3), *P*=<0.0001 (1-way RM ANOVA). Post hoc testing was performed comparing all stimulation types. *N*=136 cells; \**Q*<0.05. **g**, Schematic illustrating dorso-ventral differences in RF size, between lamina I (green) and lamina II (purple). **h**, HT RF size plotted as a function of dorso-ventral depth. **i**, Mean RF size of neurons located within lamina I and lamina II. There was a significant main effect of stimulation (*F* (1, 3) = 15.76), *P*=0.0286, and a significant interaction between stimulation and lamina (*F* (1,3) = 13.56), *P*= 0.0347 (2-way repeated measures ANOVA). Post hoc testing was performed comparing the RF size of lamina I vs lamina II for both LT (not shown) and HT stimulations. N=4 mice; \**Q*<0.05.

### CICADA: Cell-type identification by Ca^2+^-coupled activity through drug activity

A major limitation of typical population imaging studies is that, although one can visualize Ca^2+^ transients in many neurons simultaneously, it is difficult to determine the specific identity of these neurons, thereby hampering the degree to which the data can be interpreted. To overcome this limitation, we developed a novel strategy that leverages the fact that different subtypes of neurons express different combinations of GPCRs. We reasoned that pharmacological profiling could be used visualize which cells express which receptors because activation of Gq-coupled GPCRs gives rise to IP3-mediated release of Ca^2+^ from internal stores. To isolate the direct effects (cell autonomous) from indirect effects (network responses) we included the Na^+^ channel blocker tetrodotoxin (TTX) in the bath solution, which prevents action potential propagation of neuronal activity. In the presence of TTX, the cells that show a Ca^2+^ transient in response to a given agonist are only those that express its receptor, allowing us to unambiguously define cell types based on their responses to GPCR agonists applied in series at the end of each experiment (Fig. 4a).

**Fig. 4.**
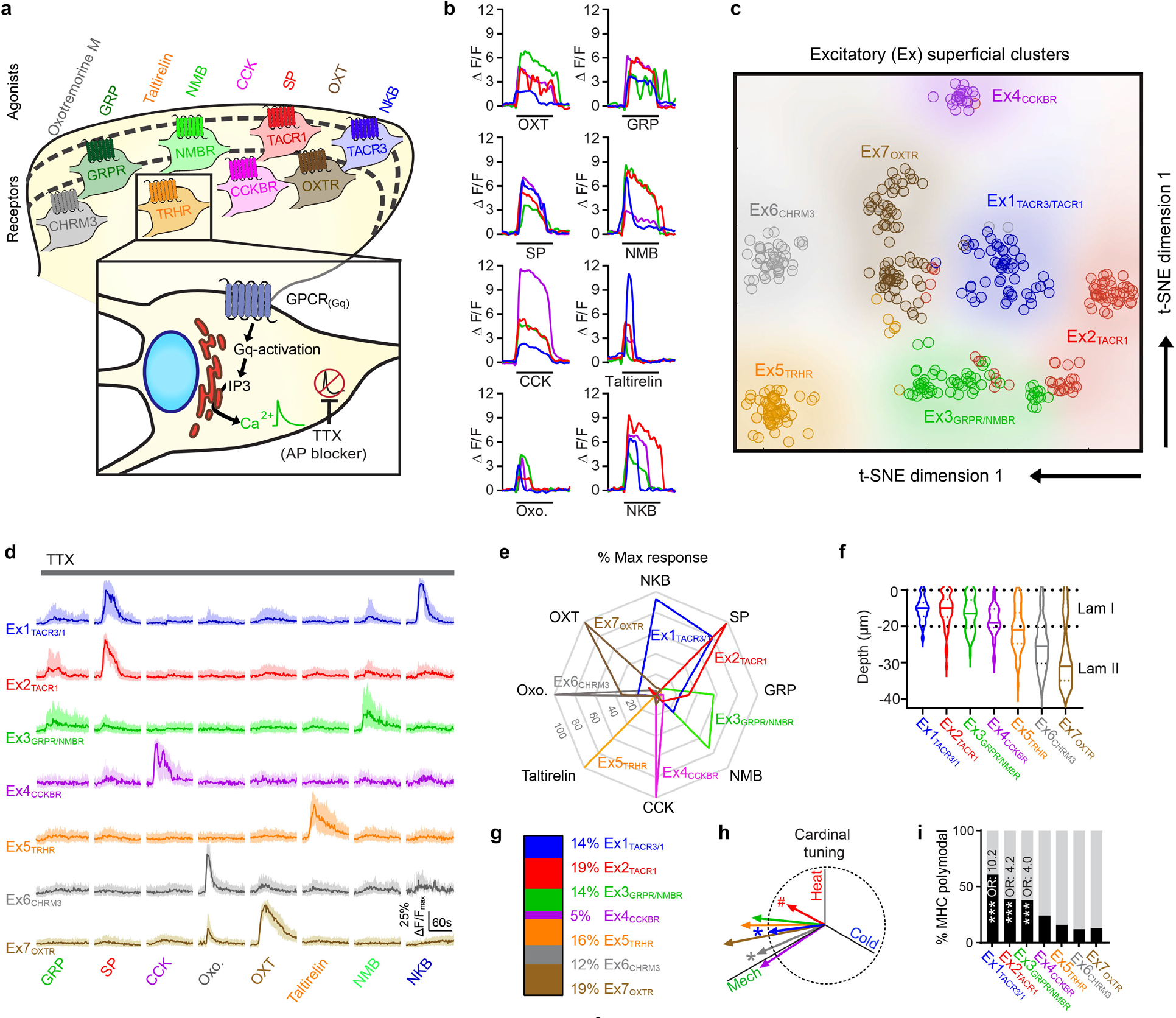
CICADA: Cell-type identification by Ca^2+^ coupled activity through drug activity. **a**, Overview of CICADA showing the intracellular mechanisms (inset) and the agonists used to target the specified receptors (top). **b**, ΔF/F Ca^2+^ traces from representative responses of individual neurons for each of the CICADA ligands. SP, 1 µM; OXT, 1 µM; GRP, 300 nM; Oxotremorine M (Oxo.) 50 µM; Taltirelin, 3 µM; NMB, 100 nM; CCK, 200 nM; NKB, 500 nM. **c**, t- SNE plot of excitatory superficial clusters (Ex) identified by CICADA. Each dot represents a CICADA-identified cell, color-coded for the appropriate cluster. Cluster numbers (*e.g.* Ex2) are followed by the principal receptor found in that population of cells (Ex2_TACR1_). **d**, Normalized Ca^2+^ traces of each CICADA ligand for Ex1-7. Each cell’s ΔF/F trace was normalized to a maximum of 1 and minimum of 0. Data are shown as median +/- interquartile range. Ex1_TACR3/1_, 57 cells; Ex2_TACR1_, 75 cells; Ex3_GRPR/NMBR_, 57 cells; Ex4_CCKBR_, 21 cells; Ex5_TRHR_, 65 cells; Ex6_CHRM3_, 49 cells; Ex7_OXTR_, 74 cells (cells are pooled from 4 mice). **e**, Radar chart showing the response to CICADA ligands within each population. The spoke length is the population average of % of maximum response to the indicated ligand. **f**, Violin plot showing the depth of each population relative to the surface of the dorsal grey matter. **g**, Relative abundance of neurons making up the excitatory populations identified by CICADA. **h**, Cardinal tuning of Ex1-7. For vector angles: 1-way ANOVA (*F* (7, 1541) = 3.983), *P*=0.0003. For vector magnitudes: 1-way ANOVA (*F* (7, 1541) = 3.289), *P*=0.0018. Post hoc testing was performed comparing each population’s vector angle (#*Q*<0.05) and magnitude (\**Q*<.05) to the average of all recorded cells. **i**, Percent of cells within each cluster which show cardinal (MHC) polymodal. Multiple logistic regression tests if slope is significantly non-zero, ***p<.0001. Odds ratio (OR) shown for significant results.

To determine which GPCR agonists would be useful for CICADA profiling, we screened 15 agonists for consistent responses in superficial excitatory neurons. This initial list represented the majority of Gq-coupled receptors with known ligands that are expressed in the dorsal horn, based on RNAseq studies [15]. Of these, eight ligands gave rise to robust Ca^2+^ transients in the superficial dorsal horn: oxotremorine M (Oxo.), an agonist for the muscarinic acetylcholine receptor M3 (CHRM3); gastrin-releasing peptide (GRP), an agonist of the gastrin-releasing peptide receptor (GRPR); taltirelin, an agonist of the thyrotropin-releasing hormone receptor (TRHR); neuromedin-B (NMB), an agonist of the neuromedin-B receptor (NMBR); cholecystokinin (CCK), an agonist of the cholecystokinin type B receptor (CCKBR); Substance-P (SP), an agonist at the Substance-P receptor (TACR1); oxytocin (OXT), an agonist at the oxytocin receptor (OXTR); and Neurokinin-B (NKB), an agonist at the tachykinin receptor 3 (TACR3) (Fig. 4b, Extended Data Fig. 5). Overall, we found that 25% of recorded neurons in the superficial dorsal horn responded to one or more CICADA ligands. Because the combinatorial responses to these ligands had the potential to be complex (Extended Data Fig. 5c-e), we next performed dimensionality reduction using K-means clustering [16]. This analysis yielded seven groups of excitatory neurons (Fig. 4c). The cluster-averaged Ca^2+^ traces revealed that most clusters were defined by a high amplitude response to one or two CICADA ligands (Fig. 4d,e). Moreover, cell clusters showed distinct distributions according to depth, consistent with the idea that they represent populations of bona fide cell types (Fig. 4f). We therefore labeled these populations Ex1 through Ex7 according to their relative location along the dorso-ventral axis and added superscript labels indicating cognate receptors. We found that the relative frequency of each population was broadly similar (∼15%), with Ex4_CCKBR_ being noticeable rarer (5%) (Fig. 4g). Notably, populations were significantly different from one another with respect to their tuning (Fig. 4h), with populations in lamina I showing a greater degree of polymodality (Fig. 4i). Thus, CICADA is a simple strategy to define populations of putative cell types in dorsal horn Ca^2+^ imaging experiments.

**Fig. 5.**
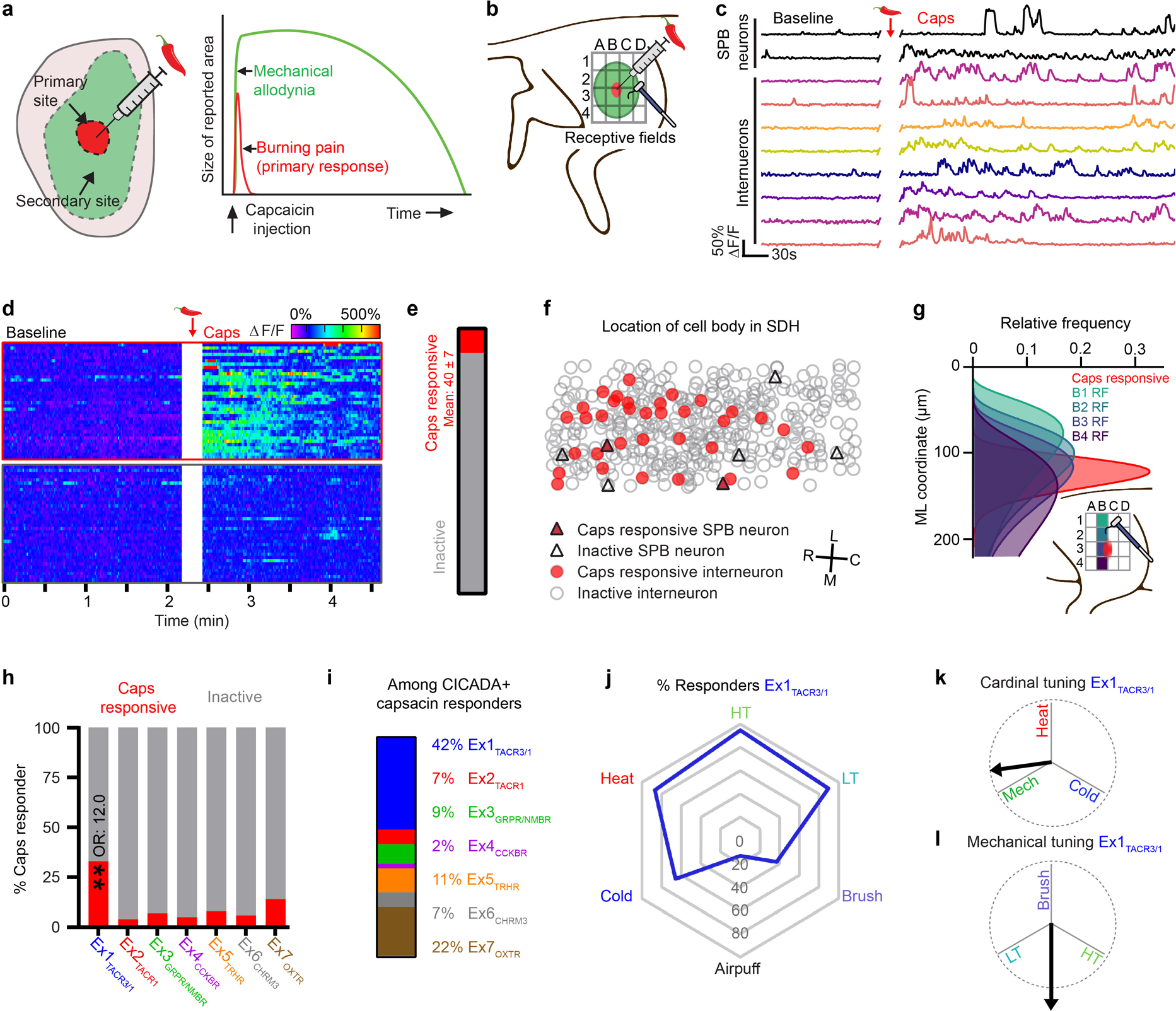
Intradermal capsaicin rapidly activates a localized subset of excitatory cells which are enriched within Ex1_TACR3/1_. **a**, Schematic of intradermal capsaicin (caps) model of central sensitization. Relative size and location of the primary and secondary sites are shown on the left and duration/intensity of the psychophysical phenomena within each region are shown on the right. **b**, Schematic for capsaicin injection and testing in the *ex vivo* preparation, the black grid shows the testing sites for receptive field mapping. **c**, Example ΔF/F Ca^2+^ traces from cells which were responsive to capsaicin injection. **d**, Heat map of ΔF/F Ca^2+^ traces from cells within the same spinal cord which were activated or quiescent after capsaicin injection. **e**, Percent of excitatory cells activated by capsaicin. N = 4 mice, mean +/- SEM. **f**, Somatic location in the SDH of SPB neurons (triangles) and interneurons (circles) which were either quiescent (open) or active (red) after capsaicin injection. **g**, Using the relative distribution of cells along the medio-lateral (ML) axis activated by HT stimulation at different receptive fields (RF) we can infer the position and proximal-distal spread of the capsaicin injection by comparing the distribution of capsaicin responders (red) to the receptive field stimulations. **h**, Percent of cells which were responded to capsaicin injection within each excitatory population. Multiple logistic regression tests if slope is significantly non-zero, **p<.001. Odds ratio (OR) shown for significant results. **i**, Relative frequency of Ex1-7 within all CICADA identified cells responsive to capsaicin. **j**, Radar chart showing the percent of cells in Ex1_TACR3/1_ which responded to the indicated cutaneous stimuli. **k**, Cardinal tuning for Ex1_TACR3/1_. (**l**) Mechanical sub-modality tuning for Ex1_TACR3/1_.

### Immediate response to intradermal capsaicin

Previous human psychophysical experiments have shown that intradermal injection of capsaicin causes an immediate response consisting of brief burning pain (<5 minutes) that is restricted to the capsaicin injection site, followed by a period of sensitization which manifests as long-lasting mechanical allodynia (several hours) extending over a large region outside the capsaicin injection site (Fig. 5a) [3]. The mechanical allodynia in this model is understood to be the result of central, rather than peripheral, sensitization because the primary afferents that innervate the secondary zone are unchanged after the capsaicin injection [5–7]. However, the specific spinal neurons and circuits involved are poorly understood.

We therefore sought to visualize how capsaicin-mediated central sensitization is manifest in the dorsal horn through Ca^2+^ imaging coupled with post-hoc cell identification through CICADA (Fig. 5b). Intradermal injection of capsaicin (7.5 µg) elicited significant increases in Ca^2+^ activity in ∼40 excitatory neurons per animal, which peaked within ∼2 minutes and persisted for ∼5 minutes (Fig. 5c-e). These capsaicin responders, which included SPB neurons, were restricted to a narrow medio-lateral band with appropriate somatotopy in the dorsal horn (Fig. 5f,g, Supplemental Fig. 2). We next analyzed the CICADA-defined populations to determine which were preferentially activated by the capsaicin injection. Using multiple logistic regression, we found that Ex1_TACR3/1_ was the only population that showed statistically significant predictive power for capsaicin response (Fig. 5h). Among capsaicin-responsive cells within Ex1-Ex7, 42% were contained within Ex1_TACR3/1_ (Fig. 5i). Neurons in Ex1_TACR3/1_ are generally polymodal (Fig. 5j) that lack strong tuning across either cardinal modalities (Fig. 5k) or mechanical sub-modalities (Fig. 5l). Thus, intradermal capsaicin predominately drives activity in Ex1_TACR3/1_, a population of relatively untuned, polymodal neurons that are located in lamina I.

### Capsaicin-induced central sensitization

Capsaicin-induced sensitization gives rise to mechanical allodynia in which LT stimuli are perceived as noxious. To interrogate the neural manifestations of allodynia at the level of the superficial dorsal horn, we compared the responses to cutaneous stimulation before and after capsaicin treatment (Fig. 6a,b). Consistent with psychophysical studies, we found that capsaicin injection altered LT responses in 56% (+/- 6% SEM) of excitatory superficial cells to LT stimuli without significantly affecting the responses to other modalities (Fig. 6c-d, Extended Data Fig. 6a,b). This change was manifest as both an increase in the proportion of neurons that respond to LT stimuli (Fig. 6e), the amplitude of response (Extended Data Fig. 6c-e), as well as increases in the receptive field size to LT stimulation (Fig. 6f, Extended Data Fig. 6f-i). Finally, analysis of mechanical sub-modality tuning revealed that, on average, neurons showed a significant shift in preference toward LT stimuli (Fig. 6g, Extended Data Fig.6j-l). To examine which cell types might be driving these capsaicin-induced changes, we compared the proportion of LT responders across CICADA-defined populations (Fig. 6h). In this analysis, three populations showed a significant increase in the proportion of cells that responded to LT stimuli post capsaicin, and were thus examined in more detail: Ex2_TACR1_, Ex5_TRHR_, and Ex6_CHRM3_.

**Fig. 6.**
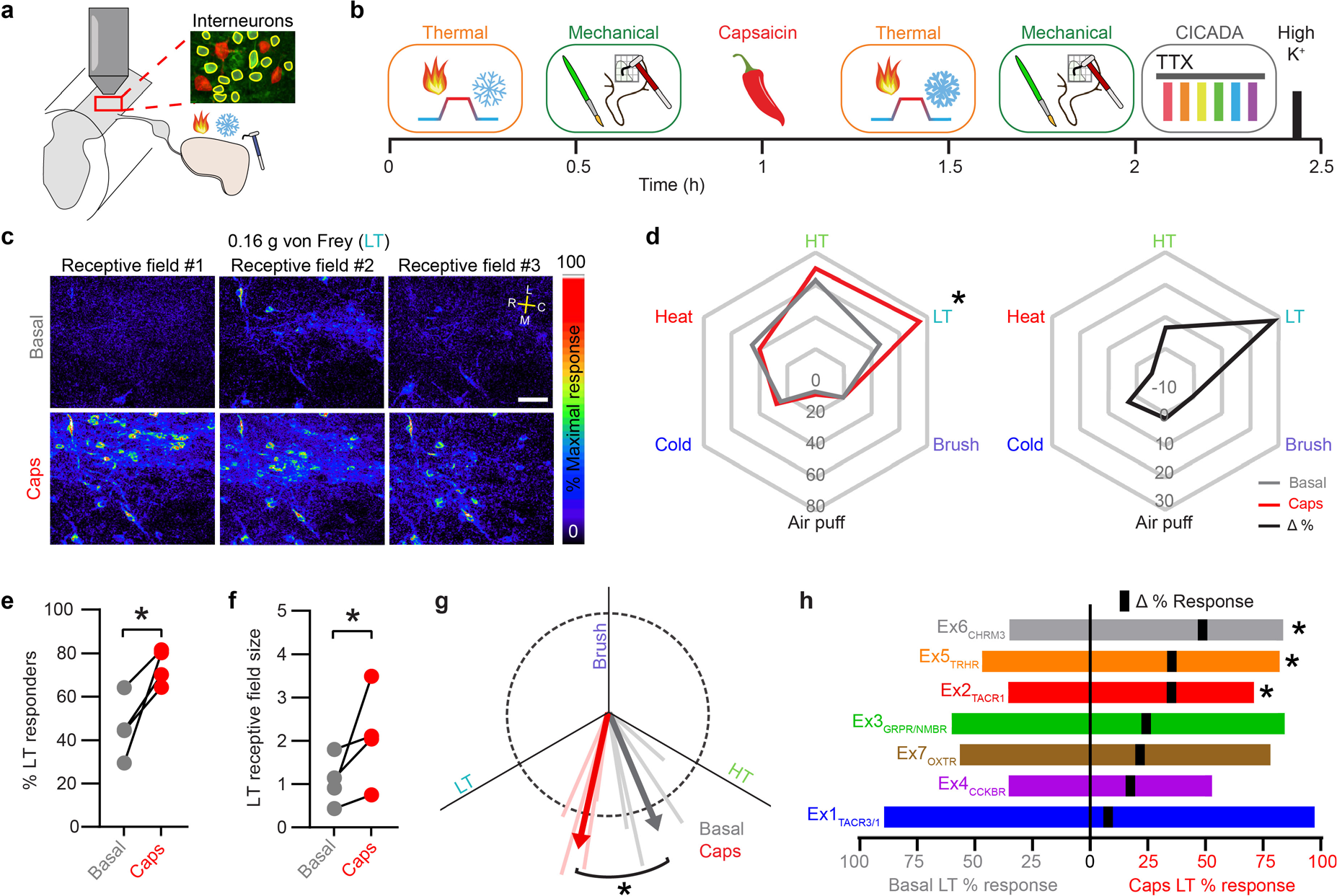
Capsaicin injection increases the frequency of LT responsive neurons and enhances LT tuning. **a**, Experimental design. **b,** Experimental protocol for testing the effects of capsaicin induced central sensitization. **c**, Field of view in the superficial SDH showing maximum ΔF/F responses during LT stimulation at 3 RFs before and after intradermal injection of capsaicin (scale bar, 50 µm). **d**, Left: radar chart showing the percent of excitatory cells responsive to the indicated stimuli before (grey) and after capsaicin (red). Right: change in percent responders. There was a significant main effect of stimulation (*F* (4, 12) = 41.11), *P*<0.0001, and a significant interaction between stimulation and time (*F* (4,12) = 5.176), *P*=0.0117 (2-way RM ANOVA). Post hoc testing was performed comparing post/pre capsaicin for all stimulations. N=4 mice, \**Q*<0.05 **e**, Percent of all excitatory cells responsive to LT stimulation before/after capsaicin. \**Q*<0.05. **f**, LT RF size of excitatory cells before/after capsaicin. Corrections for both LT and HT were included (HT data shown in Extended Data Fig. 6). There was a significant interaction between stimulation and time (*F* (1,3) = 13.34), *P*=0.0354 (2-way RM ANOVA). Post hoc testing was performed comparing RF size before/after capsaicin. N=4 mice, *Q<0.05. **g**, Mechanical sub-modality tuning for excitatory cells before/after capsaicin. Average value shown as thick arrows and individual animals shown as light, thin lines. *P>0.05 (paired t test). **h**, Percent responders to LT stimuli before (left) and after (right) capsaicin for each CICADA population. The change in % responders (Δ) is indicated with a black tick mark. There was a significant main effect of capsaicin (*F* (1, 3) = 11.02), *P*=0.0451 (RMLM model). Post hoc testing was performed comparing post/pre capsaicin for all populations, N=4 mice, *Q<0.05.

The largest capsaicin-induced change was observed in Ex6_CHRM3_ (Fig. 7a-c), a relatively untuned population found in lamina II that showed little activity in response to natural stimuli in the basal state. After injection of capsaicin, however, Ex6_CHRM3_ cells showed significantly increased responsivity across almost all modalities, including LT, HT, heat, and cold (Fig. 7a). In addition, capsaicin treatment caused a 3.9-fold increase in the size Ex6_CHRM3_ receptive fields, however, the tuning of Ex6_CHRM3_ remained unchanged (Fig. 7b,c). These findings raise the possibility that capsaicin-induced central sensitization alters the response properties of neurons in the Ex6_CHRM3_ population such that they behave as general amplifiers that increase the gain of sensory input. In addition, the increased receptive field size to LT input suggests that Ex6_CHRM3_ cells might contribute to psychophysical phenomenon that sensitization occurs beyond the original site of injury.

**Fig. 7.**
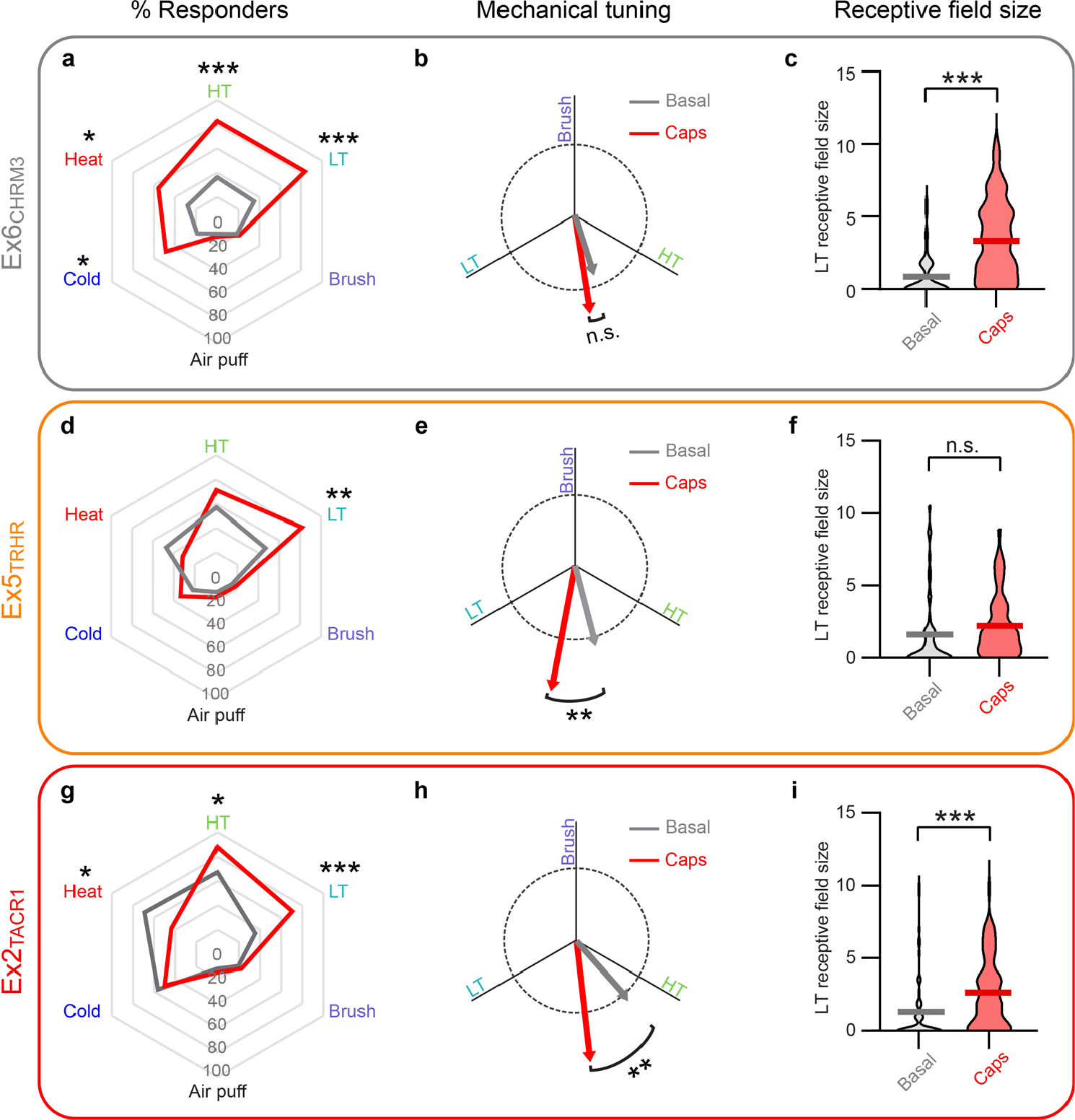
CICADA reveals distinct populations involved in central sensitization. Radar charts showing the percent of cells in Ex6_CHRM3_ (**a**), Ex5_TRHR_ (**d**), and Ex2_TACR1_ (**g**) which responded to the indicated cutaneous stimuli before (grey) and after capsaicin injection (red). *Q<0.05, **Q<0.01, ***Q<0.001, for each population a Fisher’s exact test (# responses before and after capsaicin) with FDR correction was performed for each modality, only significant results are indicated (Benjamini and Hochberg, 6 tests). Mechanical sub-modality tuning for Ex6_CHRM3_ (**b**), Ex5_TRHR_ (**e**), and Ex2_TACR1_ (**h**) before (grey) and after capsaicin injection (red). There was a significant main effect of capsaicin (F (1, 123) = 30.10), *P*<0.0001 (2-way RM-ANOVA). Post hoc testing was performed comparing post vs pre capsaicin for each group, **Q<0.01. LT RF size of Ex6_CHRM3_ (**c**), Ex5_TRHR_ (**f**), and Ex2_TACR1_ (**i**) before (grey) and after capsaicin injection (red). There was a significant main effect of capsaicin (F (1, 148) = 45.59), *P*<0.0001, and a significant interaction between capsaicin and CICADA population (F (2, 148) = 5.765), *P*=0.0039 (2-way RM-ANOVA). Post hoc testing was performed comparing post/pre capsaicin for all stimulations, ***Q<0.0001. Only cells with cutaneous input were included in these analyses. Ex2_TACR1_, 62 cells; Ex5_TRHR_, 45 cells; Ex6_CHRM3_, 43 cells (cells pooled from 4 mice). Data shown are mean of pooled cells.

The second CICADA population that showed capsaicin-induced changes to LT input was Ex5_TRHR_, another broadly tuned population found in lamina II that showed modest responses to natural stimuli in the basal state. Upon capsaicin-induced sensitization, Ex5_TRHR_ cells showed a dramatic and selective increase in their response to LT input, with no change in their response to HT, heat, or cold (Fig. 7d). This increase in response to LT was manifest as a significant shift in tuning toward LT, but no significant change in receptive field size (Fig.7e,f). Thus, Ex5_TRHR_ cells are well positioned to contribute to the psychophysical phenomenon of allodynia.

The third population that showed capsaicin-induced changes was Ex2_TACR1_, a lamina I population that is tuned for noxious stimuli under basal conditions (e.g., heat and HT, Extended Data Fig. 7). Following capsaicin treatment, however, this population showed a significant change in tuning towards LT input (Fig. 7h). These findings raised the possibility that following capsaicin treatment, the LT-induced activity in these cells may signal nociception, and thus represent a neural correlate of allodynia. In addition, capsaicin treatment significantly increased the receptive field size of Ex2_TACR1_ neurons (Fig.7i). Because the capsaicin-induced changes observed in Ex2_TACR1_ neurons appear to be a combination of the changes observed in Ex5_TRHR_ and Ex6_CHRM3_, these findings raise the possibility that Ex2_TACR1_ could be involved in integrating multiple populations to mediate allodynia within the secondary zone.

### Allodynia is manifested in spinal output neurons

In order for the sensitization we observed in excitatory interneurons to drive the psychophysical phenomenon of allodynia, spinal projection neurons must relay these nociceptive signals supraspinally. Consistent with this idea, we found that capsaicin treatment shifted the overall tuning of SPB neurons towards LT input (Fig. 8a,b). Because SPB neurons are heterogeneous with respect to their response properties, we classified them as either wide-dynamic range (WDR), high threshold (HT), or low threshold (LT) (Fig. 8c) using previously published criteria [4]. We found that both HT and WDR SPB showed significant increases in tuning to LT input, however the degree of change was much greater for WDR compared to HT SPB neurons (Fig. 8d). Finally, examining CICADA responses across populations, we found that WDR cells were more frequently responsive to CICADA agonists (71.4%), compared to either LT (40%) or HT (0%) SPB neurons. In particular, all SP-responsive SPB neurons were WDR cells (Fig. 8e). Since capsaicin is known to activate peptidergic afferents that release SP, these findings suggest that capsaicin-induced allodynia may be conveyed to the brain through a population of TACR1 expressing WDR SPB neurons and, to a lesser extent, some HT SPB neurons (Fig. 8f).

**Fig. 8.**
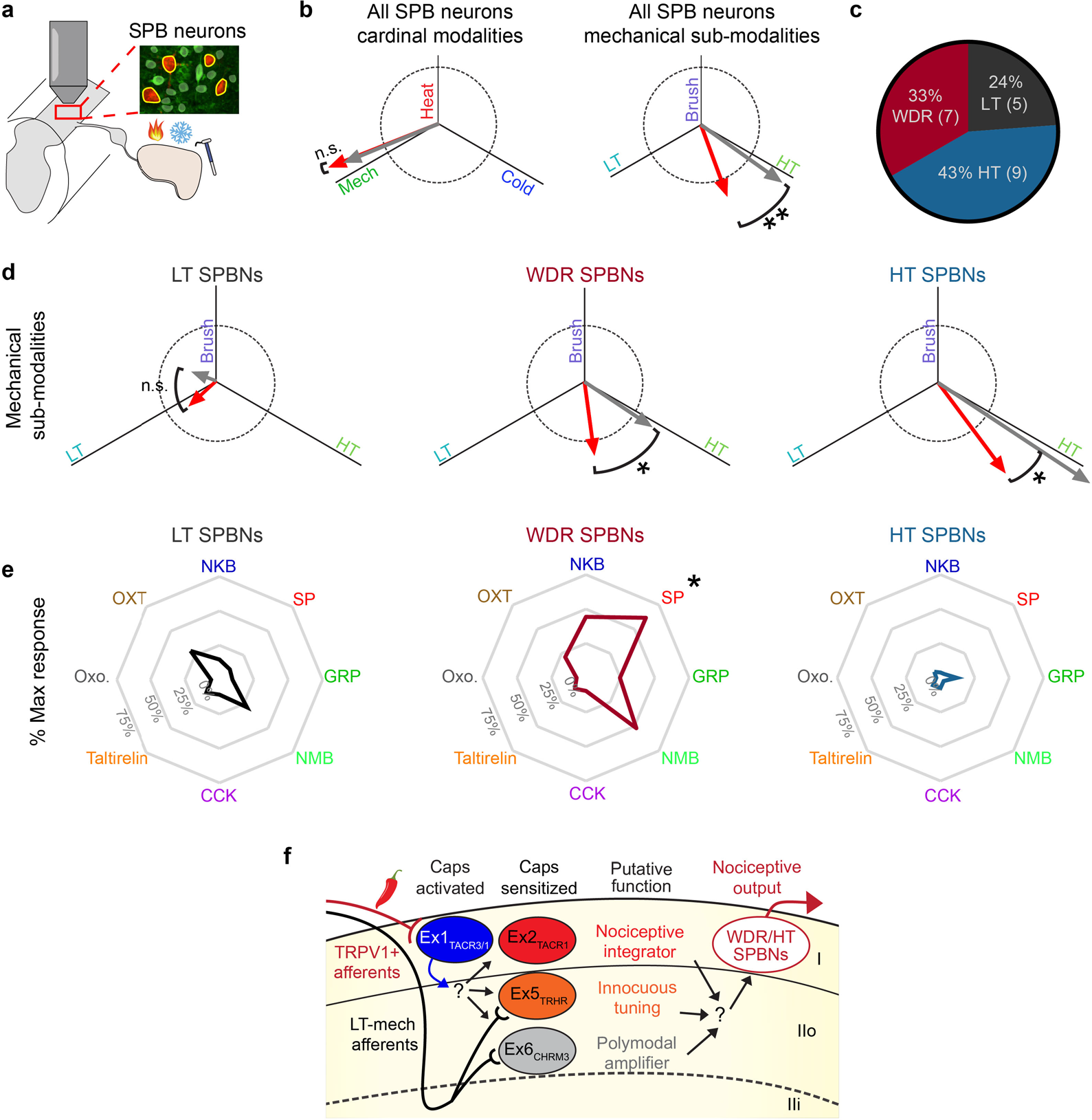
Functional classes of SPB neurons show distinct alterations in tuning after central sensitization. **a**, Experimental design. **b**, Left: cardinal modality (left) and mechanical sub-modality (right) tuning of SPB neurons (SPBNs) before (grey) and after capsaicin injection (red). **P<0.01 (Wilcoxon matched pairs signed rank test). **c**, Pie-chart of the 3 functionally defined classes within SPB neurons. Wide-dynamic range (WDR, Red) SPBNs are defined as those that responded to both low and high intensity stimuli but encoded the stimuli intensity in their response amplitude. High-threshold (HT, Blue) SPBNs responded primarily to the 2.0 g VF filament (HT stimulus). Low-Threshold (LT, Black) SPBNs only responded to brush and/or the 0.16 g VF filament (LT stimulus). **d**, Mechanical sub-modality tuning of LT (left), WDR (middle), and HT (right) SPBN sub-groups before (grey) and after capsaicin injection (red). *P<0.05 (Wilcoxon matched pairs signed rank test). **e**, Radar chart showing the response to CICADA ligands within LT (left), WDR (middle), and HT (right) SPBN sub-groups. There was a significant main effect of CICADA agonist response (*F* (3.885, 65.48) = 3.377), *P=*0.015, and a significant interaction between class and CICADA agonist response (*F* (14, 118) = 3.143), *P*=0.0003 (2- way RM ANOVA). Post hoc testing was performed comparing CICADA agonist response across functional groups, \**Q* <0.05. **f**, Proposed model and putative roles for identified excitatory populations in the development of capsaicin induced allodynia.

## Discussion

Here, we provide the first comprehensive functional characterization of neurons in the dorsal spinal cord at a population level. First we applied quantitative natural stimuli to the skin to determine response properties of individual neurons. To further classify these neurons, we developed CICADA, a novel method to identify neuronal subsets based on their responses to Gq-coupled GPCR ligands. We then describe the spinal representations of transient burning pain and persistent mechanical allodynia that arise from intradermal injection of capsaicin. Our findings suggest that different CICADA-defined populations contribute to distinct aspects of the capsaicin response. In particular, Ex1_TACR3/1_ neurons are immediately responsive to capsaicin injection, whereas Ex2_TACR1_, Ex5_TRHR_, and Ex6_CHRM3_ differentially contribute to the spinal representation of allodynia. Finally, we see altered tuning of SPB neurons, which shift towards LT stimuli, thereby providing a neural substrate through which normally innocuous information could be conveyed to the brain via a nociceptive output pathway.

### Ex1_TACR3/1_ neurons initiate central sensitization

The application of capsaicin to the skin activates TRPV1-expressing primary afferents, which target the most superficial aspect of the dorsal horn and release the neuropeptides SP and CGRP. Our study revealed that Ex1**_TACR3/1_** neurons are the primary neuronal population in the dorsal horn that responds to this afferent input. Importantly, the time course of capsaicin-induced activity in Ex1**_TACR3/1_** neurons matches the duration of acute pain that follows capsaicin treatment. Compared to neurons in other excitatory populations, Ex1**_TACR3/1_** neurons are the most superficial, the most polymodal, and have the largest receptive field sizes (Extended Data Fig. 7). These findings suggest that Ex1**_TACR3/1_** neurons are likely involved in mediating capsaicin-induced pain and initiating central sensitization.

### Central sensitization involves distinct neuronal elements

We found that three CICADA- defined populations showed capsaicin-induced alterations in their response properties to low threshold mechano-sensory input and, strikingly, these neural populations changed in different ways. Neurons in the Ex6_CHRM3_ population showed a dramatic increase in receptive field size coupled with amplified responses to most types of sensory stimuli, but no overall change in tuning. In contrast, neurons in the Ex5_TRHR_ population showed a selective increase in their responsiveness to LT mechanosensory input, with no change in receptive field size. Finally, neurons in the Ex2_TACR1_ population showed a combination of these two features, with both an increase in receptive field size and a shift in tuning towards LT input. These capsaicin-induced changes in activity are consistent with a model in which distinct lamina II populations mediate separate aspects of capsaicin-induced sensitization—amplification and tuning—which are integrated by interneurons in lamina I before being conveyed to the brain (Fig. 8f). Although this model remains somewhat speculative because we do not yet have a detailed wiring diagram of the dorsal horn, it may nevertheless serve as a starting point to study the neural circuitry of central sensitization.

### Neural correlates of psychophysical phenomena

In psychophysical studies of capsaicin-induced central sensitization, participants report two salient phenomena: touch stimuli evoke pain (allodynia), and the region of sensitivity extends beyond the initial site of injury (extended secondary zone). To help visualize the neural basis of allodynia, we developed a vector-based method to represent neuronal tuning to somatosensory stimuli. We found that capsaicin treatment alters the tuning of several neuronal populations such that they are now almost equally responsive to LT and HT inputs. Because these neurons were tuned for noxious input under basal conditions, this altered tuning is suggestive of a neural correlate of allodynia. To help visualize the neural basis that might account for an extended secondary zone, we developed an approach for population-level receptive field mapping. We found that two CICADA-defined populations showed an increase in receptive field size following capsaicin treatment. Thus, the overall changes in neural activity described here may account for two of the most salient features of capsaicin-induced sensitization. Human psychophysical experiments have revealed that heat and brush responses are likewise sensitized by capsaicin, although these effects are more transient and occur over a smaller area of skin compared to punctate mechanical allodynia [3]. A limitation of our study is that we did not capture the corresponding neural representations of thermal hyperalgesia or dynamic allodynia, likely because the Peltier thermode and brush stimulus used in this study were applied to the entire receptive field, thereby limiting our ability to see localized changes in the dorsal horn.

### Towards the modeling of somatosensory integration

Because prolonged activity in TRPV1- positive afferents gives rise to the release of SP and CGRP in the dorsal horn, it is tempting to speculate that these neuropeptides may be involved capsaicin-mediated sensitization. In this regard, it is noteworthy that many of the neurons that showed altered activity following capsaicin (Ex1**_TACR3/1_**, Ex2**_TACR1,_** and the WDR SPB neuron subtype) express the receptor for SP. Whether any of these populations also express receptors for CGRP remains unclear. The receptor for CGRP is Gs-coupled (rather than Gq-coupled) and, as a result, the activation of this receptor is not visible using the current CICADA strategy. The future development of tools and strategies that allow us to visualize Gs- and Gi-coupled signaling events would not only aid in our understanding of neuromodulatory mechanisms, but also permit a much more complete analysis of neuronal subtypes through pharmacological profiling. The combination of population imaging and pharmacological profiling described here are the first steps of an approach that will enable new the modeling of somatosensory integration at a new level.

## Methods

### Animals

All experiments were performed with approval of the University of Pittsburgh’s IACUC. Mice that were heterozygous for both the *Vglut2-ires-cre* allele (Jax Stock #016963) the *Ai96* allele (for Cre-dependent expression of GCaMP6s, Jax Stock # 028866) were used for all experiments. Mice were housed on a 12-hour light cycle with *ad-libitum* water and food in accordance with the United States National Institutes of Health guidelines for the care and use of laboratory animals. Ca^2+^ imaging was performed on mice of both sexes ranging from 5-10 weeks of age.

### Parabrachial dye injections

Mice ranging from 4 – 5-weeks old were placed under isoflurane anesthesia and mounted in a Kopf stereotaxic headframe. After exposing and leveling the skull a burr hole was made at (RC/AP: -5.2 mm from Bregma or -1.5 mm; Interaural line ML: 1.25 mm; DV: 3.0 mm from bregma or 2.23 mm from skull) and a glass micro-pipette filled with Fast DiI oil (2.5 mg/mL; Invitrogen, Carlsbad, CA) was used to inject 250 nl into the left parabrachial nucleus over the course of 5 min. The pipette was left in place for 10 min before retracting and wound closure. Injected mice were then used for imaging a minimum of 4 - 5 days after injection. DiI-labeled SPB neurons were validated by taking fresh transverse spinal cord slices and coronal brain sections and checking for fluorescence in the appropriate regions as mapped in a reference atlas.

### Semi-intact somatosensory preparation

Dissections were performed described previously [17] with some noted differences to account for optical recordings. In brief, mice were deeply anesthetized with a ketamine/xylazine mixture (1.75 mg ketamine and 0.25 mg xylazine per 20 g body weight). The right hind paw, leg, and back were shaved with an electric clipper leaving 2 - 3 mm of hair in place. Next, mice were transcardially perfused with 95%O_2_/5%CO_2_ saturated, chilled, sucrose-based artificial cerebrospinal fluid (ACSF). The sucrose-based ACSF contained (in mM): 234 sucrose, 2.5 KCl, 0.5 CaCl_2_, 10 MgSO_4_, 1.25 NaH_2_PO_4_, 26 NaHCO_3_, and 11 Glucose. A dorsal laminectomy was performed followed by complete excision of the spinal column, right ribs, and right leg. The tissue was transferred into a dissection dish and the lateral femoral cutaneous and saphenous nerves were dissected out in continuum from the skin to the DRG. The thoracolumbar spinal cord was then removed from the bone and the roots were carefully pulled away from the cord to allow movement of the spinal cord. The spinal cord was pinned at an angle such that the grey matter of the spinal cord was parallel with the ground. The dura mater was removed among the entire spinal cord and the pia was only removed along the L1 - L2 recording area. The roots and nerves were loosened from the tissues as much as possible and the skin was pinned out as far away from the spinal cord as possible. This ensured sufficient separation from the spinal cord as to allow a 3D printed light shield to block the visible light used to localize stimulation on the skin from reaching the high sensitivity detectors on the microscope. Once the skin and spinal cord were securely pinned in place using Minutien pins (FST Cat: 26002-20) the dish was transferred to the microscope and washed with standard normal ACSF solution containing (in mM): 117 NaCl, 3.6 KCl, 2.5 CaCl_2_, 1.2 MgCl_2_, 1.2 NaH_2_PO_4_, 25 NaHCO_3_, 11 glucose), which was slowly washed in over 15 min. Next, the temperature of the wash solution was raised from room temperature to 28°C in 2°C steps every 5 - 10 min. Once the tissue had equilibrated to the standard ACSF, a manual wash was performed to fully remove any leftover detritus and dissection solution using a 50 ml syringe filled with ACSF. Imaging began 30 - 45 min after the final wash in order to allow time for thermal expansion and settling of the tissue. Once ∼1 L of solution was washed through the dish, the remaining 4 L of bubbled ACSF was then recirculated for the duration of the physiology experiments.

### Multiphoton Imaging

Ca^2+^ imaging was performed on a Leica SP5 MP microscope using a Leica 20x NA 1.0 water immersion lens with a Coherent Chameleon Ultra II Ti:Si femtosecond laser set to 940 nm for Ca^2+^ imaging and 1040 nm for imaging of DiI within the SPBPNs. Fluorescence was captured with HyD detectors and standard FITC/TRITC emission filter sets with a 570 nm dichroic beam splitter. Three different planes of the superficial grey matter were simultaneously imaged with a spacing of 14 µm between planes, which yielded a range of depths from 0 to 60 µm below the spinal cord surface. For each plane, scanning was performed at 1 Hz with a 1.4x optical zoom at 512 X 214 pixel resolution and a field size of 528.2 x 220.8 µm size yielding a pixel resolution of 1.03 µm/pixel. This volumetric scanning allowed sampling from multiple lamina simultaneous and typically yielded ∼300 excitatory neurons in a given experiment.

As the location of afferent input for each animal is determined by relative somatotopy, the region of the grey matter imaged thus needed to be functionally identified for each animal. The exact field of view imaged in the SDH was determined by using a low magnification image and a search stimulus within the dissected skin. A wide brush which stimulated the entire length of the skin was used to activate as many afferent inputs as possible. The imaging field of view was iteratively refined, centering the imaging on the region that has the most change in fluorescence and then zooming in until a final zoom was reached. Subsequently a 2.0 g von Frey (VF) filament was used to delineate a 15 x 15 mm region of skin was delineated that showed the most input into the final field of view in the SDH. The region was marked with a surgical felt tip pen. Imaging, stimuli controllers, feedback sensors (thermocouples, load cells, etc.), and event times were synchronized using a Power1401 and Signal (CED).

### Drug Applications

After the final round of sensory testing post capsaicin was completed, 500 nM tetrodotoxin (TTX) was applied and allowed to circulate for 15 - 20 minutes before proceeding. CICADA agonists were then applied in a randomized order for 2 min followed by 5 min wash in the continued presence of 500 nM TTX. CICADA agonists were kept frozen in single use aliquots until application when they were thawed and diluted in TTX recording solution. Post CICADA, a brief pulse of 5 ml of ACSF containing 30 mM KCl was injected into the perfusion lines in order to confirm cell viability and assist in ROI generation and refinement. All drugs were purchased from Tocris and Sigma.

### Determination of dorso-ventral location

In order to accurately calculate the depth of cells in the dorsal horn, the curved surface of the dorsal horn needed to be taken into account. Briefly, after all experimental manipulations had finished, a volumetric time-lapse (XYZT) imaging protocol was loaded at the same field of view and XY resolution as the Ca^2+^ imaging protocol but encompassing the entire recorded volume of the grey matter at high Z resolution taken from ∼10 µm above the surface to ∼60 µm below, at a Z resolution of 1 µm. During this time, 30 mM KCl was applied in order to visualize all the GCaMP6s expressing cells and provide a high contrast image for volumetric reconstruction. Post-hoc, the structural reference was aligned with the functional Ca^2+^ imaging reference image, smoothed with a 3D median filter, contrast enhanced using Contrast Limited Adaptive Histogram Equalization, and then XY down sampled into voxels representing a 10 x 10 x 1 XYZ area. The mean value of the voxels was then used to find the surface of the dorsal horn grey matter for each 10 x 10 XY coordinate. The calculated Z coordinate of the surface was based on identifying the asymptotic increase in fluorescence as the surface of the dorsal horn comes into focus. Each of the 3 recording planes was then aligned to a Z-plane in the structural image and each ROI was then offset in the Z-dimension by the amount indicated by their XY location. This analysis provided the final dorso-ventral location of each cell.

### Natural cutaneous stimuli application

The presentation order of natural stimuli was randomized within each testing block (i.e. baseline or post capsaicin testing periods). All natural stimuli were applied to an identical 15×15 mm area of skin using stimulators that were specifically designed to fit in the same footprint. A dim red light was focused on the skin in order to provide sufficient illumination for the experimenter and reduce the amount of light contamination reaching the sensor. For manually applied stimuli, a program was written in Signal (CED) to produce a synchronized output to an audio monitor that provided a 3,2,1 countdown for each stimulus, a 1s tone for duration of application, and ensured both even spacing and duration of stimuli application.

### Mechanical Stimuli

Airpuff was applied at a constant supply pressure of 60 PSI for 1 s duration and repeated 3 times 60 s apart and controlled by a PicoSpritzer III (Parker Hannifin). The airpuff applicator device was a custom designed module with a 6 x 6 grid of evenly spaced 18-gauge stainless steel nozzles with an inner diameter of 0.8 mm to fit within the delineated region (15 x 15 mm) and provide equal pressure across the skin. The device held in place 1 cm above the skin with a micromanipulator. Brushing was done manually with a quantitative sensory testing brush (Somedic SenseLab AB, SENSELab Brush-05). Average force applied was ∼20 g for brushing and ∼5g for airpuff and was calibrated by use of a load sensor coupled to the imaging chamber and controlled by an Arduino. As brushing was manual, we did observe some variance in the force applied but it typically stayed within the range of 15 to 25 g. The applied pressure for each mechanical stimulus was estimated as follows: air puff, 0.2 kPa; brush, 6.5 kPa; 0.16 g VF (LT), 78 kPa; 2.0 g VF (HT), 270 kPa. While the Somedic brush is a 15 x 5 mm brush with an area of 75 mm^2^, we estimated the thickness of the brush when compressed during a stroke was ∼2 mm resulting in an effective area closer to 30 mm^2^ when actively applying pressure. The brushing was repeated 3 times, 60 s apart.

Receptive field mapping was done with a LT (0.16 g) or HT (2.0 g) von Frey filament. A custom cut flexible plastic grid was placed upon the delineated region of skin in order to assist with consistent targeting of the filaments to each of the 16 sites to be mapped in a 4 x 4 grid arrangement. Each round of mapping consisted of applying the filament for 1 s to each location spaced 15 s apart. For example, in Fig. 3D, the first region stimulated was A1 then B1, C1, D1, A2, B2, *et cetera*. Once the final location was reached the mapping was repeated once more for a total of 2 replicates for each filament and testing block.

### Thermal stimulus application

A custom-made water-cooled Peltier was placed on the delineated region of skin and held at a thermoneutral 30°C between stimulations. The Peltier was then driven by a custom-designed Peltier controller to the indicated temperatures at specified rates and set point. Measured surface temperatures were recorded in Signal with a CED 1401. For fast rate heat ramps the Peltier was driven at 4°C/s from 30°C to 52 °C and held for 5 s at 52°C before returning to neutral. Fast cold ramps were similarly run at 4°C/s but the effective cooling rate slows as it approaches 0°C (as seen in Fig. 1G). Thermal steps were held for 15 s at the intermediate temperature (40°C for heating and 18°C for cooling) and stepped to the subsequent temperature (52°C or 4°C) at maximum ramp rate and held for 5 s once at the final temp. Both heating and cooling slow ramps run at a constant rate of 1°C/s from either 30°C to 52°C or 30°C to 4°C. All thermal stimuli were applied twice for each testing block separated by at least 1 min.

### Capsaicin Injection

After the first testing block was completed, 7.5 µg of capsaicin dissolved in 10 µl of PBS containing 0.5% Tween-80 was intradermally injected with a 31-gauge needle to the result of a visible bleb. Because additional light was needed for consistent needle placement, imaging was stopped during this time to prevent PMT damage, resulting in ∼30 s blind spot around the injection time. The next block of sensory testing was performed 5 min following injection, followed by repeated blocks of testing ∼30 min apart. The effects of capsaicin on mechanical activity were seen at least 15 min after capsaicin injection and remained consistent for the 3 rounds of post capsaicin testing performed. For analysis here, the middle block of capsaicin testing (∼30 min post injection) was used for all animals.

### Image processing and data extraction

For image processing, we utilized a Suite2p pipeline empirically tuned for superficial dorsal horn cells for image registration and initial segmentation. The masks were transferred to ImageJ and additional ROIs were added by hand as the heterogeneity in morphology and size in the superficial layers of the spinal cord was difficult for most segmentation programs to accurately capture. Masks were then spot checked using a maximum activity (ΔF/F or the first derivative of fluorescence across time) reference image generated from 5-min bins along the entire recording. This maximum activity reference image resulted in ∼50 frames that represented 5- min bins of activity and could be visually scrolled through to quickly check the 4- to 5-h long recording for physical deformations, drift, blebbing, or other pathologic changes. Any cells that showed significant deviation from their XY location, signs of Z-drift (e.g., changes in the appearance of the nucleus, soma, or processes), blebbing, or cell death in the latter portions of the recording were manually deleted. Once the masks were finalized, the mean fluorescence was calculated and any cell that has significant drift (>200%) in their 5-min binned median fluorescence value was deleted because such a change was indicative of either physical drift or Ca^2+^ dysregulation which would signal signs of apoptosis. Then ΔF/F was calculated for each cell using a rolling ball baseline: for each frame (F_i_) the 30^th^ percentile value of the surrounding 5 minutes of frames was used as a baseline (F_b_) to perform the ΔF/F calculation ((F_i_- F_b_)/F_b_). Rolling ball normalization was preferred here as the effects of slow photobleaching of the indicator and/or small changes in location were minimized in these long recordings.

### Analysis of Ca^2+^ activity

Once ΔF/F values had been generated, traces were analyzed for binary responder analysis (yes/no) as determined by a peak value in ΔF/F that was both >125% of baseline in the 5 s following stimulation and also > 6σ (standard deviations) of basal trace noise in the 30 s prior. These criteria were selected based on a receiver operator characteristic (ROC) analysis of σ and amplitude using a manually annotated data set of 500 cells as the ‘ground truth’ against which we optimized the AUC from the ROC analysis. For the manual annotation, a Matlab script was used to generate image files for each cell with stimulation event markers. The 500 traces were then manually marked as a responder or not based on whether the amplitude of response was clearly distinguishable over the background recording noise. This ‘ground truth’ data was then used to calibrate against multiple objective measures of quantification (Extended Data Fig. 3). We chose to slightly favor specificity over sensitivity for these analyses to minimize false positives. The responses to CICADA agonists used an identical amplitude and σ criteria but with a 120 s window of response to match the duration of application. This window also accommodates the higher variation in onset of response due to the pharmacokinetics of drug penetration and variations in intracellular signaling cascades of the different agonists. Euler diagrams were made using eulerAPE [18].

Once the activity had been binarized, responses to the various stimuli were assigned. For brush and airpuff stimulation, a cell was considered a responder if it met threshold criteria for at least 2 of 3 applications to be considered a responder. For heating, cooling, or static LT/HT stimuli if a cell responded to at least 1 of the stimuli it was considered a response. This was due to the fact that each of the heating and cooling ramps have distinct parameters (*e.g.*, fast vs slow rate of change) and the static LT and HT stimulation each stimulate a distinct region of skin so each of the stimulations were not equal. To calculate the size of the receptive field, the total sum of responses at the 16 tested fields (4 x 4 grid) were averaged across 2 trials providing a possible receptive field size of 0 to 16. As each of the 16 tested sites were spread across a 225 mm^2^ region spaced apart by ∼3.75 mm, each response likely represents ∼3-4 mm^2^ of skin, *i.e.* a cell which responded to 4 points has an estimated receptive field size of 15 mm^2^. For clarity, the conversion to square mm^2^ was not performed and was left as number of receptive fields.

For capsaicin injection, a cell was considered responsive to capsaicin if the activity post capsaicin injection was greater than 150% of pre-capsaicin levels based on a 2-min average of ΔF/F values.

### Vector Analysis

Vectors were calculated using the maximum amplitude of response across all trials for the indicated modality. For each cell, the 3 modalities to be compared by vector analysis were first normalized to the average response. The angle of the vector was assigned 120° apart from each other modality as follows: heat/brush (+90), mechanical/LT Static (+210) and cold/HT static (+330). Using these angles and the normalized amplitudes, 3 separate vectors were calculated, summed, and then graphed in Matlab where the length of the vector was the normalized amplitude of response, and the direction of each vector was as described for each modality. For examples in interpreting these vectors we can examine Cells #1-3 from (Fig. 2g, more examples in Supplemental Fig. 1). Cell #1 has a very clear and strong tuning for mechanical stimuli. The magnitude and angle of the bi-modal mechanical and heat responsive Cell #2 indicates a tuning for heat over mechanical stimuli. In contrast the trimodal Cell #3 shows no tuning. Cells which did not have any responses prior to capsaicin were not included in the vector analysis.

### K-Means clustering

K-Means clustering was performed in the statistical package Orange[16]. Normalized amplitudes of CICADA responses were used as an input for the K-means calculation and the data was visualized using t-Distributed Stochastic Neighbor Embedding (t-SNE)[19].

### Statistical Analysis

Data are expressed as the mean ± SEM unless otherwise indicated in figure legend. The following tests were used for statistical analyses: Student *t* test (paired where appropriate *e.g.* before and after capsaicin) for comparison of exactly 2 groups, ordinary 1-way analysis of variance (ANOVA) with multiple comparisons post hoc test if indicated by the main effect (comparison of >2 groups), 2-way repeated measures ANOVA (restricted maximum likelihood (REML) was used where balanced repeated measures data was unavailable) with multiple comparisons post hoc test if indicated by the main effect (time course with comparison of 2 or more groups). All tests were 2 tailed and a value of *P*<0.05 was considered statistically significant in all cases. For cases where multiple post hoc comparisons were indicated by the main effect, a correction for multiple comparisons was implemented using a False Discovery Rate of 0.05 via the two-stage step-up method of Benjamini, Krieger and Yekutieli. Reported *Q*- values are the FDR corrected *P*-values, and in all cases *Q*<0.05 was considered significant. For all data plotted as a function of depth (*e.g.* Fig. 3h) smoothing was applied using Prism (9 neighbor, 6^th^ order polynomial). All other statistical analyses were performed using GraphPad Prism 9 software.

## Acknowledgements

The research reported in this publication was supported by the National Institute of Neurological Disorder and Stroke of the National Institutes of Health under Award Number R01 NS096705 to H.R. Koerber, NS073548 to C. Warwick, and the National Institute of Arthritis and Musculoskeletal and Skin Diseases of the National Institutes of Health under Award Number R01AR063772 to S.E. Ross.

## Author Contributions

Author contributions are based on CRediT taxonomy to ensure clarity. (https://credit.niso.org/)

| Contributor Role | CAW | JS | JH | KMS | HC | JI | HRK | SER |
| --- | --- | --- | --- | --- | --- | --- | --- | --- |
| Conceptualization | X |  |  |  |  |  | X | X |
| Data Curation | X | X |  |  | X |  |  |  |
| Formal Analysis | X | X |  |  | X | X |  |  |
| Funding Acquisition |  |  |  |  |  |  | X | X |
| Investigation | X | X | X | X |  |  |  |  |
| Methodology | X | X |  |  |  | X | X | X |
| Project Administration | X |  |  |  |  |  | X | X |
| Resources | X | X | X | X |  | X |  |  |
| Software | X | X |  |  |  |  |  |  |
| Supervision |  |  |  |  |  |  | X | X |
| Validation | X | X |  |  |  |  | X | X |
| Visualization | X |  |  | X |  |  |  |  |
| Writing – Original Draft Preparation | X |  |  |  |  |  |  |  |
| Writing – Review & Editing | X | X | X | X | X | X | X | X |

## Extended Data Figure Legends

**Extended Data Fig. 1.**
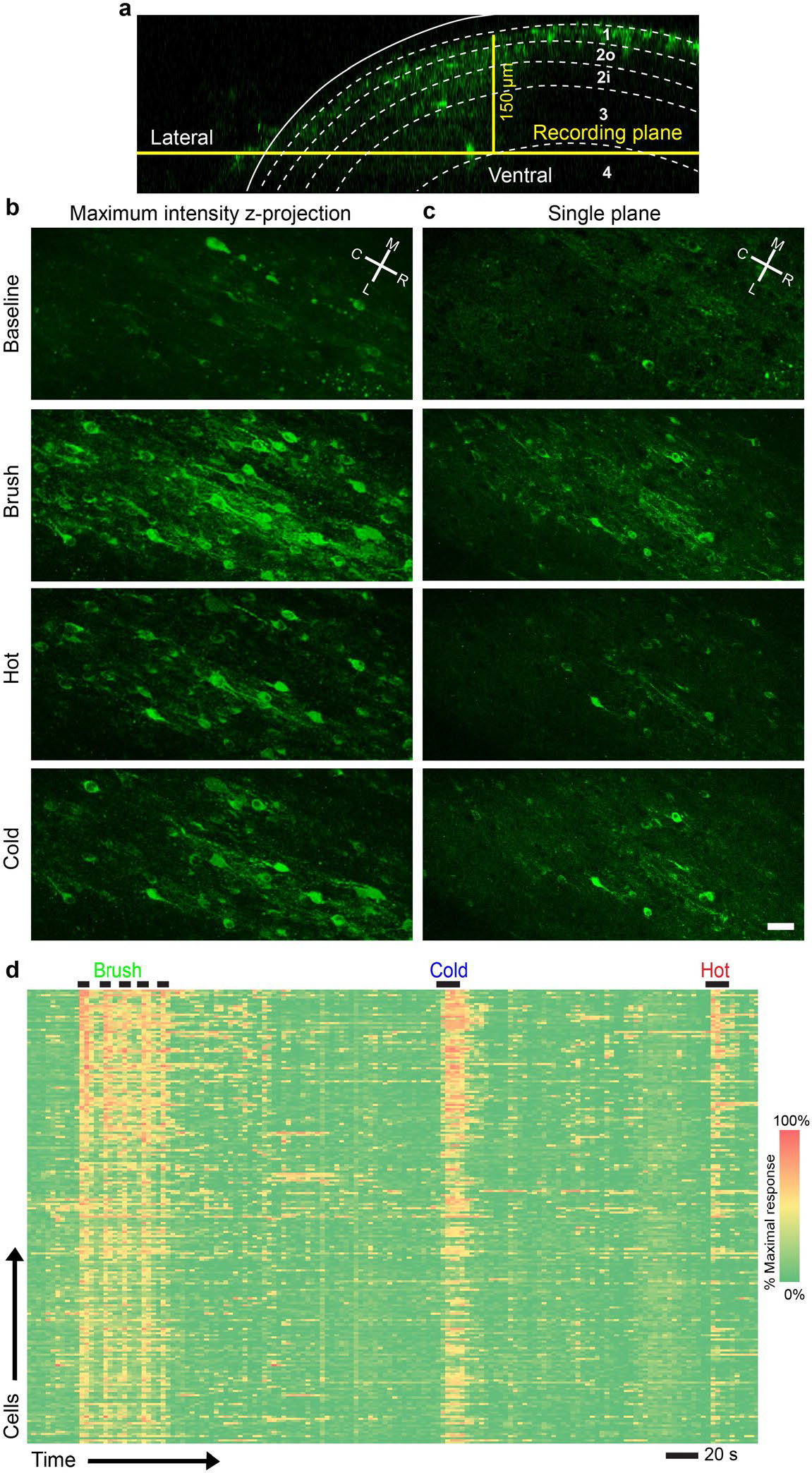
Imaging cells beyond Lamina II provides similar signal to noise. **a**, GCaMP6s expression in *Vglut2-Cre* positive excitatory neurons in an orthogonal view showing a reconstructed XYZ image (*i.e.* a transverse slice) taken post imaging showing the recording position relative to the surface of the dorsal horn. **b**, Maximum intensity Z-projection of 3 image planes separated by 10 µm during brush, hot saline, or cold saline application. **c**, Individual imaging plane highlighted in **a** (scale bar, 30 µm). **d**, Heat map of Ca^2+^ responses in deeper lamina cell to brush, cold saline, or hot saline applied to the skin.

**Extended Data Fig. 2.**
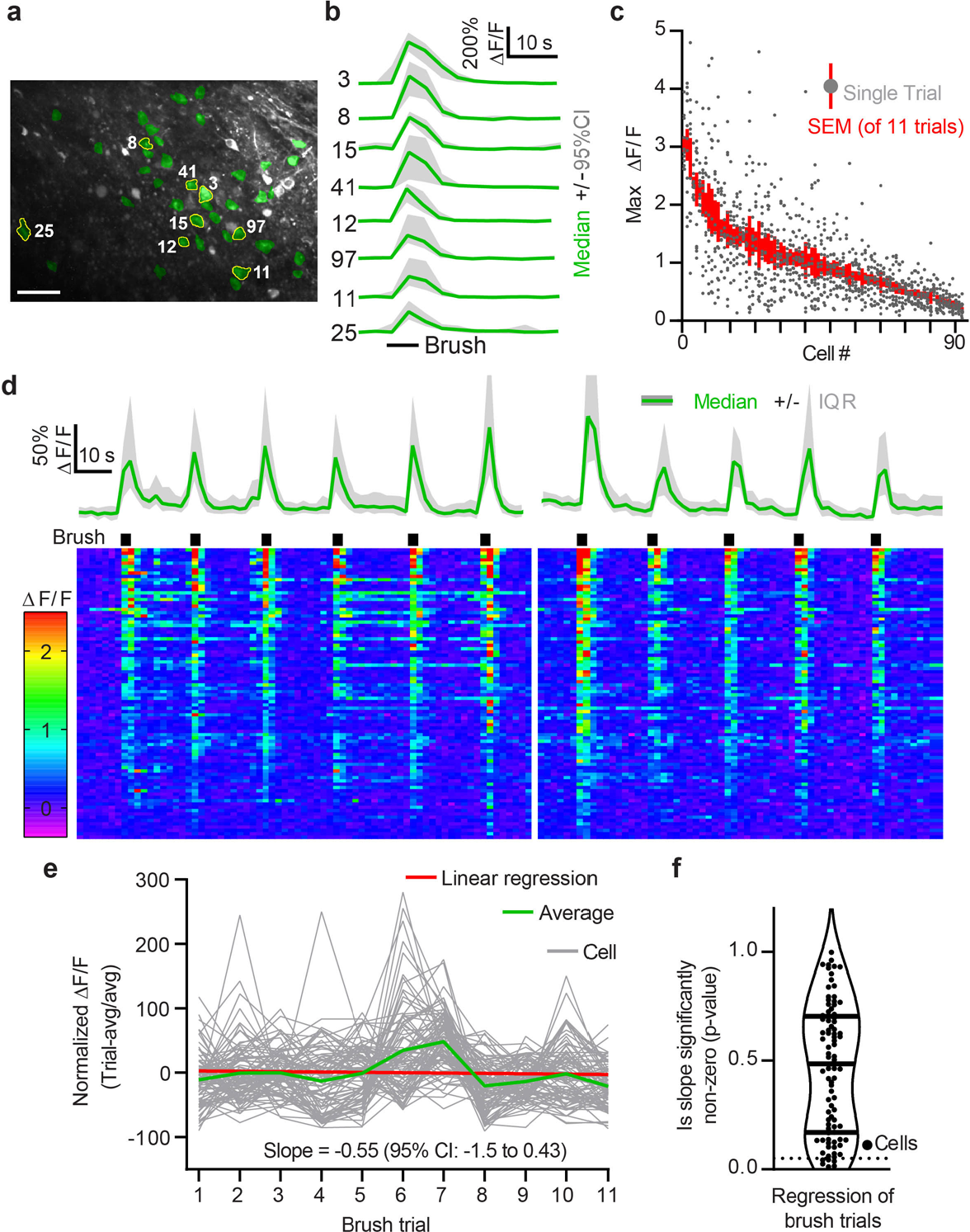
Repeated cutaneous stimulation does not produce rundown of response. **a**, Superficial (<50 µm) GCaMP6s expression in *Vglut2-Cre* positive excitatory neurons. Brush responsive cells are highlighted in green (scale bar, 50 µm). **b**, ΔF/F Ca^2+^ traces from the circled cells in **a**. Median (green) +/- 95% CI (grey) showing the consistency within 11 brush trials. **c**, ΔF/F Ca^2+^ amplitudes in response to brush in one example field of view. **d**, Top, ΔF/F Ca^2+^ traces median (green) +/- interquartile range (grey) of brush responsive cells. Bottom, heatmap of ΔF/F Ca^2+^ traces. **e**, Linear regression analysis of brush responses shows no correlation between the brush trial and the normalized response across the average of all cells. **f**, *P*-values of regression analysis to test whether the slope of each individual cell’s brush responses are non-zero. The vast majority of individual cells (dots) have no significant trends in their responses.

**Extended Data Fig. 3.**
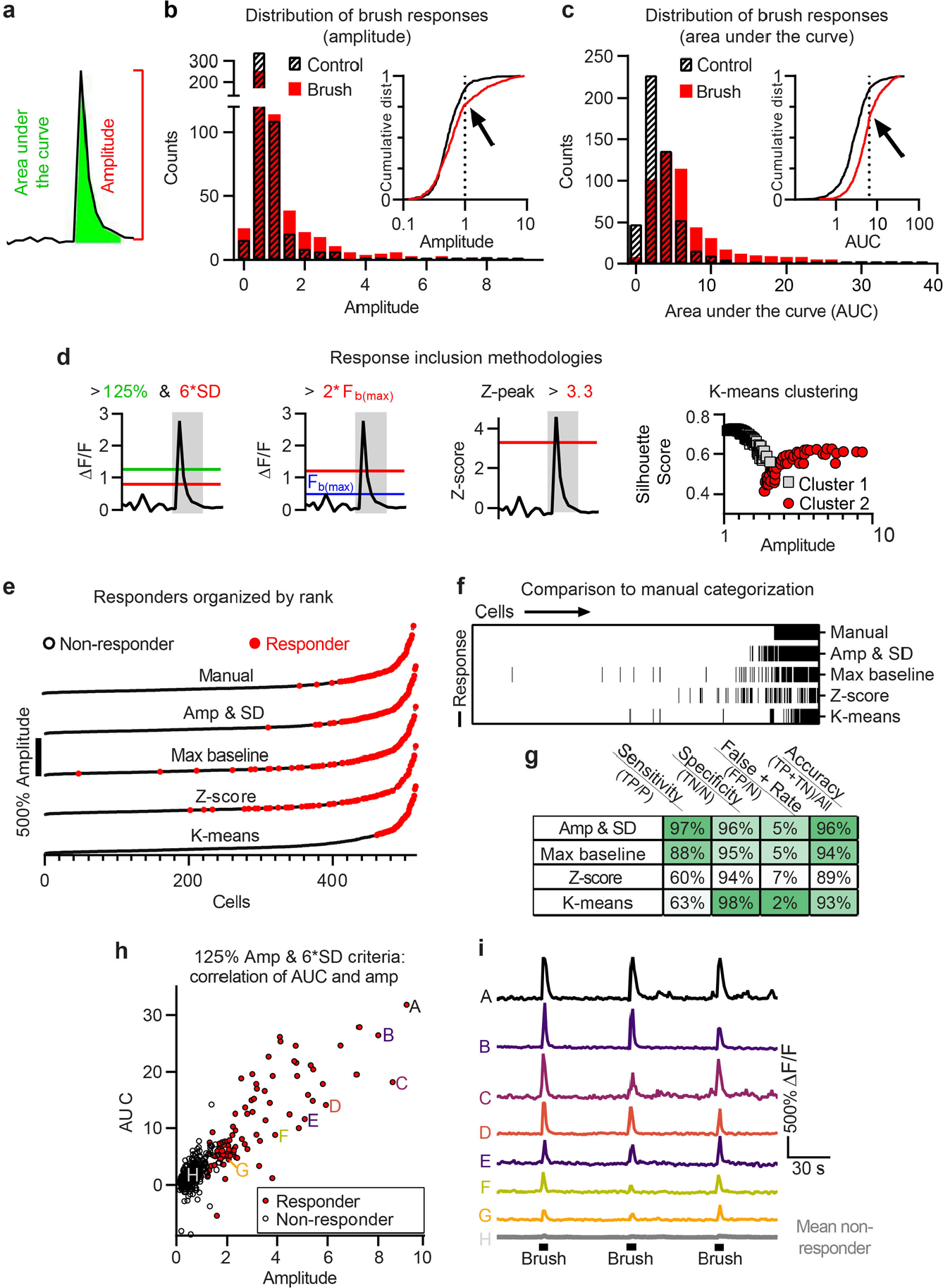
Determination of threshold of response. **a**, Schematic of quantification metrics for Ca^2+^ signals. **b**,**c** Histogram of amplitude and AUC values from brush stimulation or an equivalent period of time immediately prior to stimulation. Inset: cumulative distribution of each value showing the inflection point at which two populations are readily distinguishable. **d**, Schematic of other published methods for determination of a ‘responsive’ neuron. **e**, Responses to brush organized by ranked amplitude response. Each line is the same set of cells determined as a responder (red) or non-responder (black) by the indicated method. **f**, Responses to brush ordered by manual categorization and then by amplitude. Black ticks indicate a responsive cell according to the indicated method. **g**, Comparison of different methods relative to manual review. True positive (TP), positive (P), true negative (TN), false positive (FP), negative (N). In general, we found that using a two-factor determination (*i.e.* Amplitude and SD) provided the best balance of the available methods. Methods like K-means had an exceptional specificity but very low sensitivity which could be preferable in some scenarios. Using the maximum fluorescence at baseline and also provided very good overall accuracy at the expense of a noticeably smaller sensitivity. This is due to the high penalty incurred by having a small amount of spontaneous activity at baseline, which is generally not seen in DRG where the method was originally utilized [20]. **h**, Scatterplot of amplitude vs AUC with 125% & 6 SD criteria overlaid in red. **i**, Individual ΔF/F Ca^2+^ traces for labeled cells in **h**.

**Extended Data Fig. 4.**
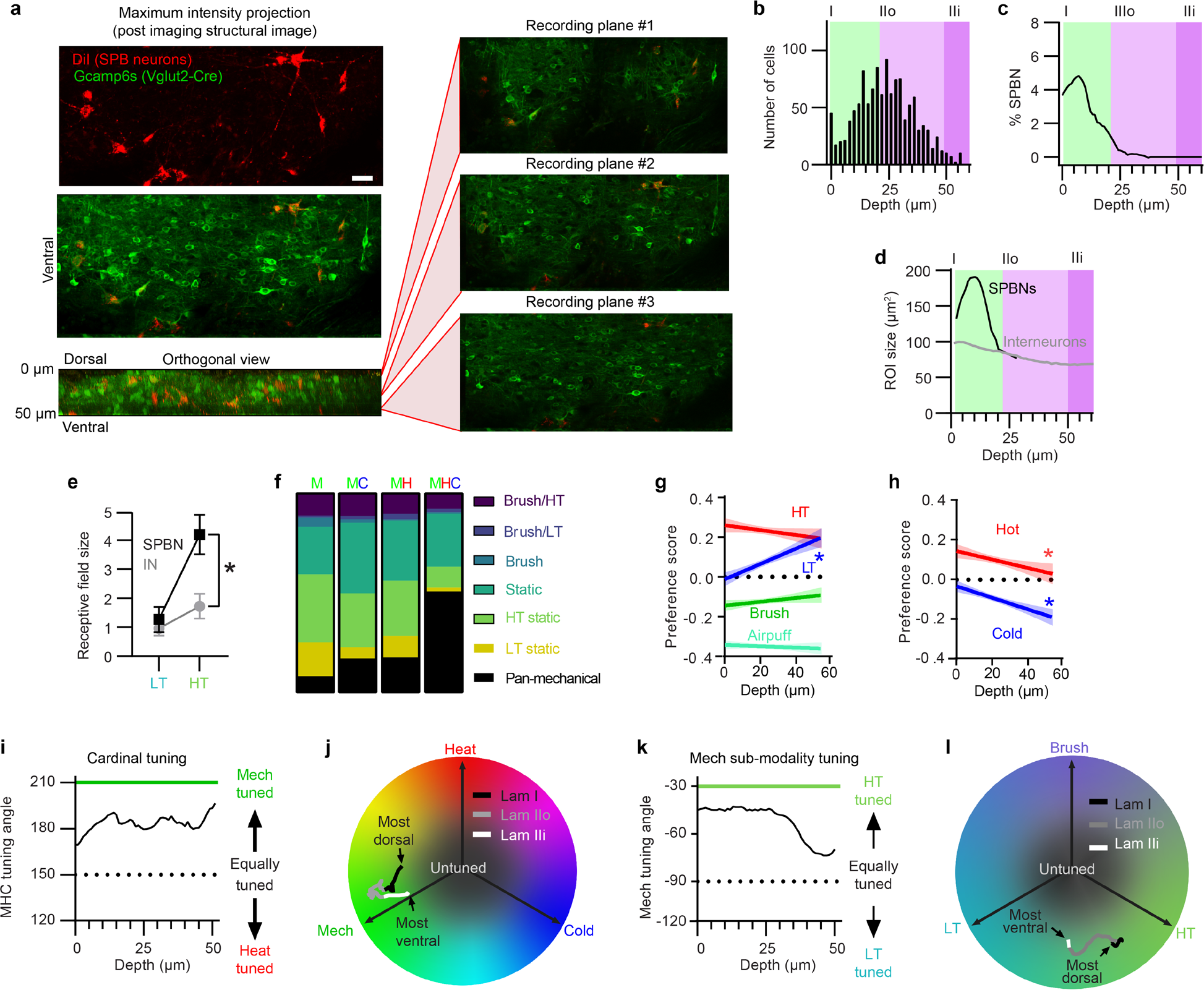
SDH DV distributions affect sensory tuning. **a**, Example images of a typical field of view within the SDH showing GCaMP6s expression in *Vglut2-Cre* positive excitatory interneurons (IN, green only) and spinoparabrachial neurons (SPBN, red and green). Maximum intensity projection images (left) are shown of the 3 recording planes (right). Images are taken at the end of each experiment in the presence of a high potassium solution to visualize viable cell bodies (scale bar, 40 µm). **b**, Histogram of the depth of sampled neurons relative to the surface of the grey matter. **c**, % of cells at each depth which were SPB neurons (*i.e.* labeled by DiI). **d**, Average region of interest (ROI) size of SPBNs and INs at each recorded depth, as a surrogate for cell body size. **e**, Receptive field sizes for LT (0.16 g filament) and HT (2.0 g filament) stimuli in SPBNs and INs. SPBNs had a significantly larger RF for HT stimuli compared to INs. There was a significant main effect of stimuli type (*F* (1, 3) = 273.7), *P=*0.0005, and a significant interaction between stimuli and neuron type (*F* (1, 3) = 1, 3), *P*=0.0392 (2-way RM ANOVA). *Q <0.05. **f**, Breakdown of mechanical sensitivity by cardinal modality sensitivity. Note that in general, the more polymodal cells also respond to more types of mechanical sub-modalities, *e.g.* MHC polymodal cells are >50% pan-mechanical vs <10% pan-mechanical for bi-modal polymodality. **g**,**h** Linear regression of preference score vs depth with 95% C.I. error bars, \**P* <0.05. Consistent with prior literature, thermal stimuli are more preferred in the superficial layers whereas among the mechanical sub-modalities only the LT stimulus shows any significant correlation to depth. Preference score is the average modality amplitude subtracted from an individual modality amplitude and then the product is divided by the maximum amplitude of response: (X_i_-X_avg_)/X_max_. **i**, Cardinal tuning angle as a function of depth shows a similar concept to preference score with only the most superficial cells (<10 µm) showing significant tuning for heat. **j**, Cardinal tuning angle and vector amplitude plotted across Lam I (black), Lam IIo (grey) and Lam IIi (white) from most to least dorsal. This plot indicates that moving ventrally in the SDH causes changes in both tuning angle as well as the magnitude. In particular, the most superficial cells are more tuned for heat, Lam IIo cells tend to have high amplitude responses (high magnitude vector), whereas Lam IIi begins to show smaller magnitude mechanical responses relative to other modalities all while following the general trend of preferring mechanical stimuli at deeper lamina. **k**, Mechanical sub-modality tuning angle as a function of depth showing that the linear regression in LT preference seen in **h** is primarily occurring at the boundary of Lam IIo and below. **l**, Mechanical tuning angle and amplitude plotted across Lam I (black), Lam IIo (grey) and Lam IIi (white)from most to least dorsal. In contrast to the cardinal modalities mechanical sub-modalities tend to just shift preference (angle) and less so magnitude potentially indicating the convergence of HT and LT input in these layers.

**Extended Data Fig. 5.**
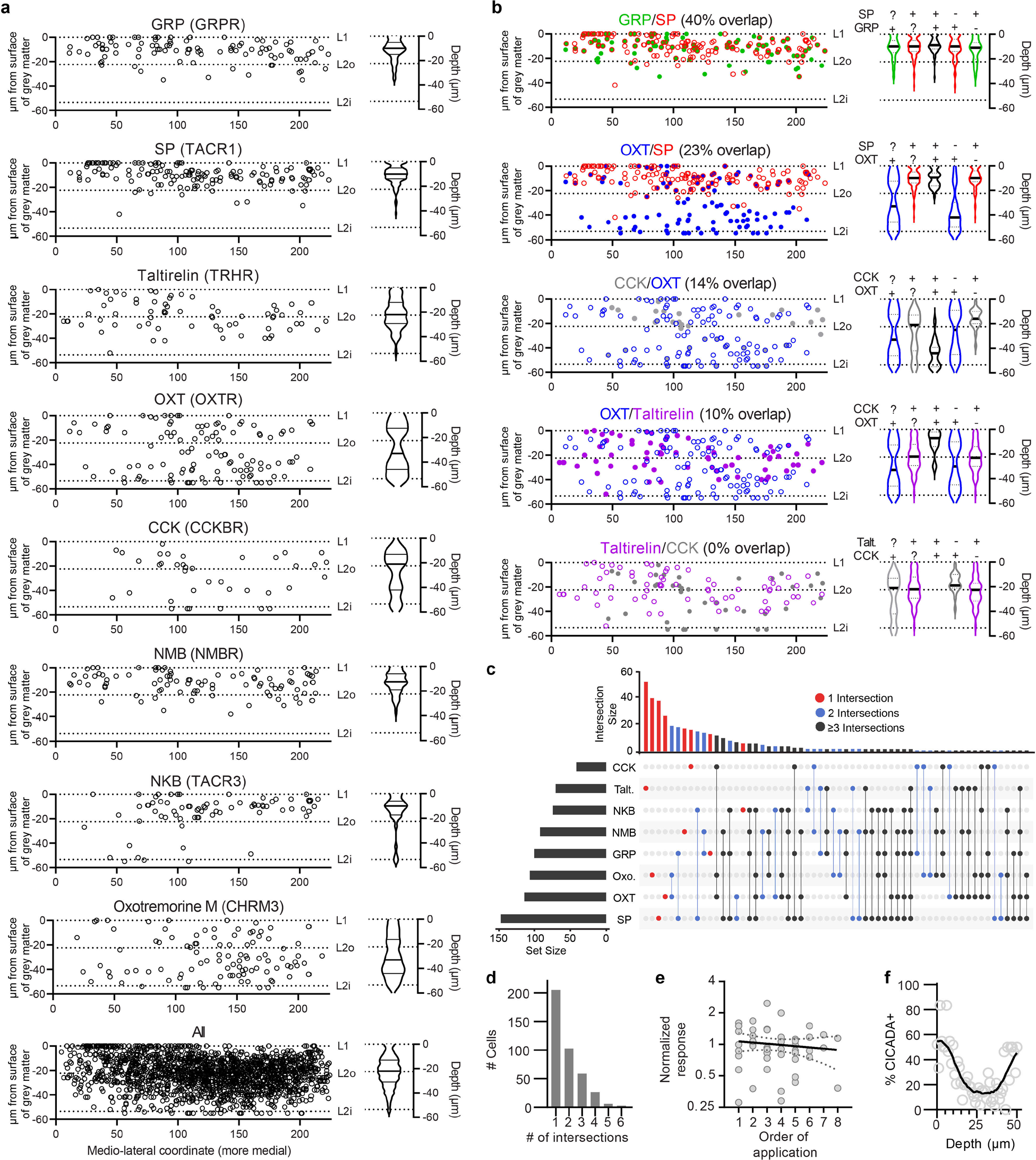
Characterization of CICADA Ligands. **a**, Dorso-ventral and medio-lateral distribution of CICADA ligand responses. Individual cells are shown on the left and violin plot of the dorso-ventral distribution on the right. **b**, Assessment of overlap between CICADA ligand responders, individual cells on left and violin plots showing the distribution of the indicated population. For violin plot, ‘+’ indicates a required ligand response, ‘- ‘ indicates no response and ‘?’ indicates either response is included. **c**, UpSetR plots for all CICADA ligands quantifying the degree of overlap/intersection between populations, 1 response (red), 2 (blue), and ≥3 (black). **d**, Histogram of ligand co-responsiveness (intersections) from the UpSetR plot in **c**. **e**, Linear regression analysis of normalized ligand response against order of application. No statistically significant effect of application order was found (DFn, DFd (1,45), Y = -0.02656*X + 1.096, *P*=0.42). Data taken from 8 animals with a total of 47 randomized applications. **f**, Percent of cells which responded to 1 or more of the CICADA ligands by depth. The average of individual 1 µm bins are shown with open circles and the smoothed average is shown with a black line.

**Extended Data Fig. 6.**
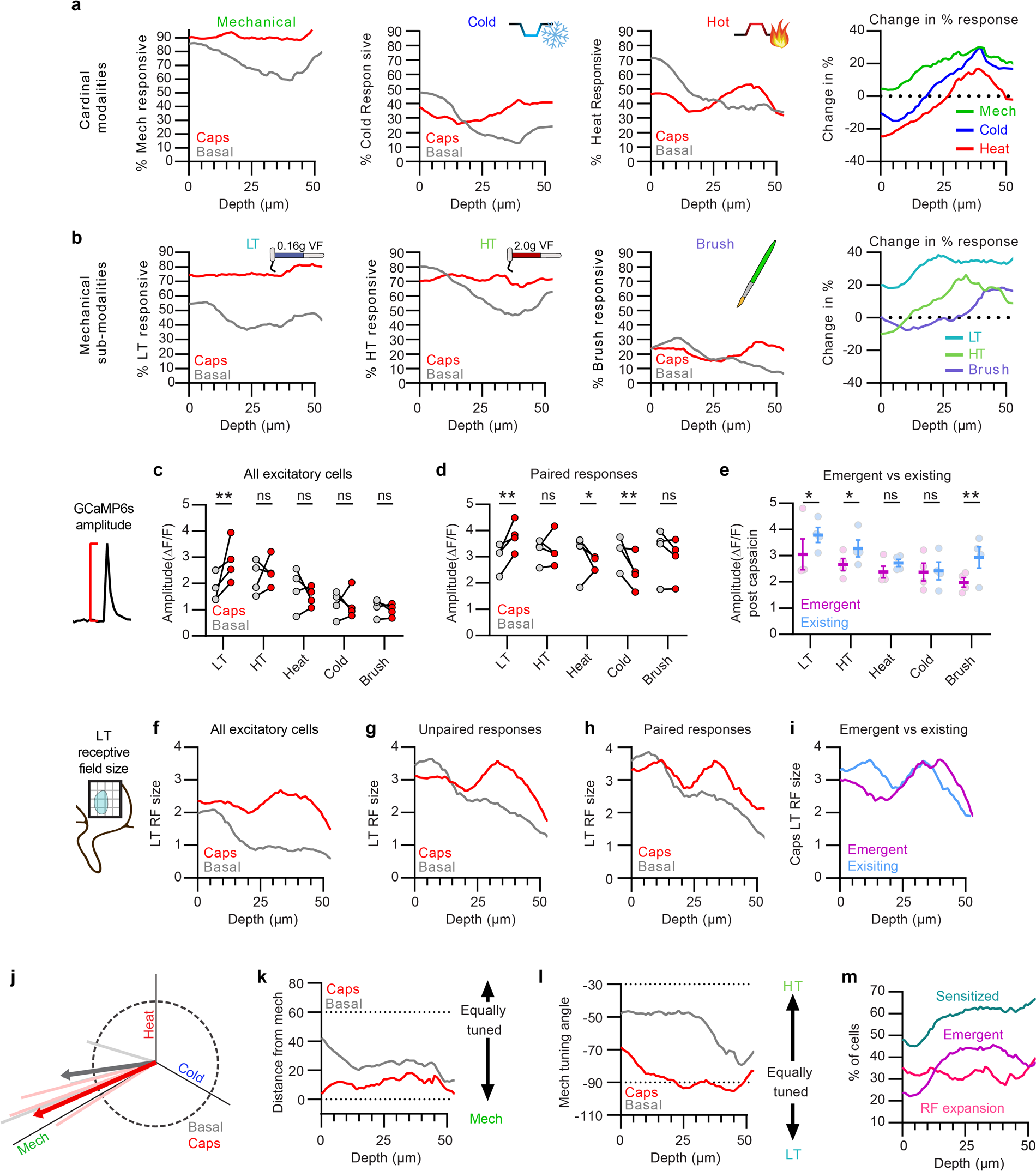
Alterations in SDH responsiveness after capsaicin vary by depth and basal response1. **a**,**b**, Percent of cells responsive to cardinal (**a**) or mechanical sub-modalities (**b**) before (grey) and after capsaicin (red) across depth. Change in % response on right. **c**, Amplitude among all excitatory cells before/after capsaicin. There was a significant main effect of stimuli type (*F* (4, 12) = 58.14), *P*<0.0001, and a significant interaction between stimuli and Caps (*F* (4, 12) = 6.522), *P*=0.005 (2-way RM ANOVA). **d**, Amplitude among cells which responded to the indicated stimulus both before and after capsaicin (paired responses). There was a significant main effect of stimuli type (*F* (4, 12) = 5.003), *P*=0.0132, and a significant interaction between stimuli and Caps (*F* (4, 12) = 11.06), *P*=0.0005 (2-way RM ANOVA). **e**, Amplitude after capsaicin among cells which were basally sensitive (blue) versus those which had an emergent response post capsaicin (purple). There was a significant main effect of stimuli type (*F* (4, 12) = 9.017), *P*=0.013 (2-way RM ANOVA). **f-h**, LT receptive field sizes before/after capsaicin within: all excitatory neurons (**f**), only LT responsive (**g**), only paired LT responses *i.e.* a response before and after capsaicin (**h**). **i**, LT receptive field size post capsaicin in emergent vs existing populations. **j**, Cardinal tuning of all excitatory neurons. **k**, Cardinal tuning across depth before/after capsaicin. **l**, Mechanical sub-modality tuning across depth before/after capsaicin. **m**, Summary of LT changes post capsaicin. Sensitized cells include cells which had at least one of the following: an emergent response, a receptive field expansion >0.5, or those with a >1.5-fold increase in amplitude. For all tests, \**Q* <0.05. N=4 mice, for **c-e** each dot represents the average of all cells within that mouse before or after capsaicin. All other graphs represent the pooled data from 4 mice with a total of 1,265 cells.

**Extended Data Fig. 7.**
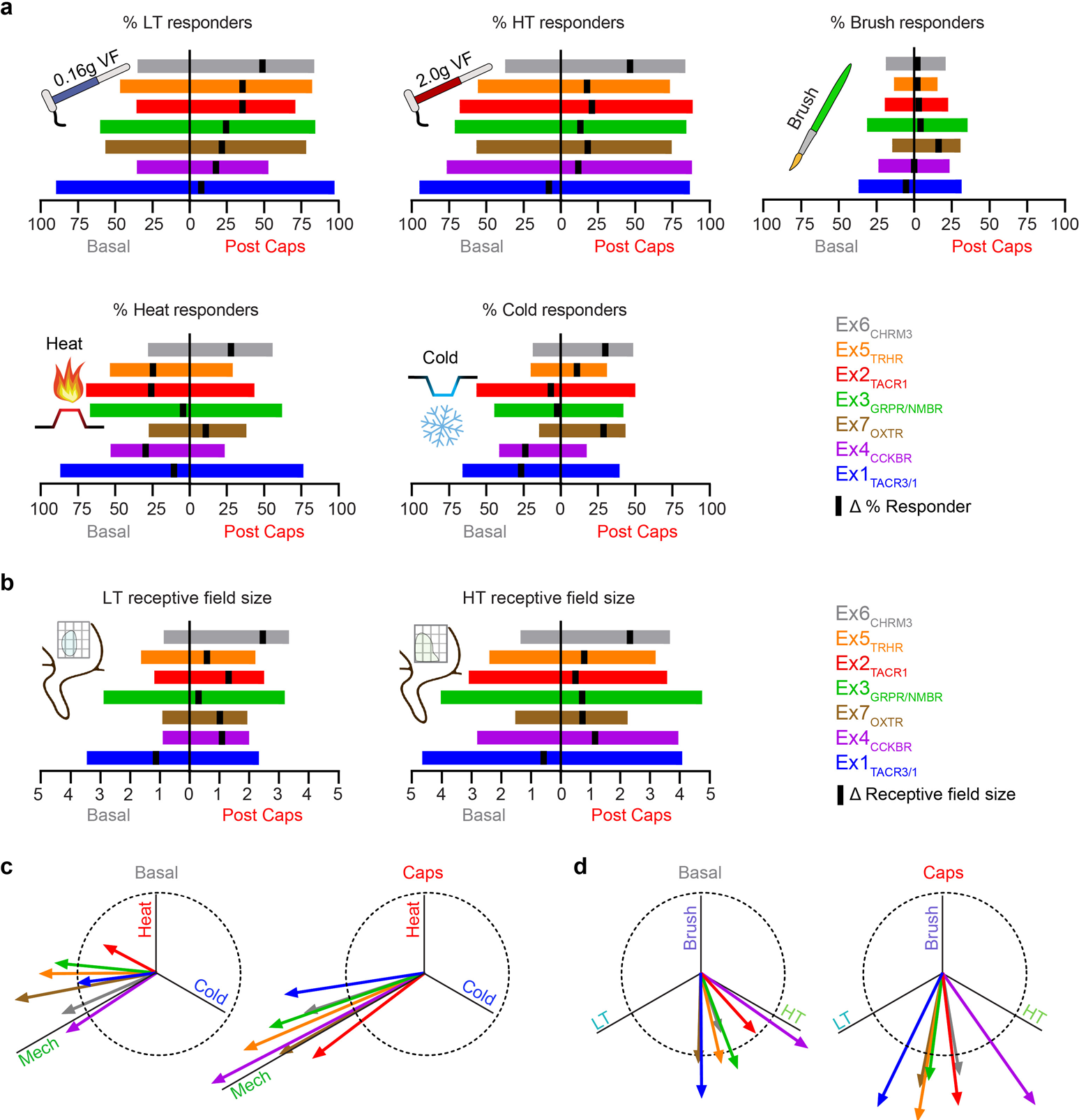
CICADA Populations show distinct basal response properties and alterations after capsaicin. **a**, Percent of cells responsive to the indicated modality before (left of y-axis) and after capsaicin treatment (right of y-axis) with the change in % responders (Δ) indicated with a black tick on the right axis for an increase in the percent of cells responsive and on the left axis for a decrease. CICADA populations are color coded and arranged in order of high to low change in low-threshold (LT) percent responders. Bars are the mean value of 4 animals. **b**, Receptive field sizes for low-threshold and high-threshold (HT) stimulations before/after capsaicin with the change (Δ) in RF size indicated by a black tick mark where increases in RF size are on the right and decreases on the left. CICADA populations are arranged as described in **a**. Bars are the mean value of 4 animals. **c**, Cardinal tuning of Ex1-7 before (left) and after capsaicin (right). **d**, Mechanical sub-modality tuning of Ex1-7 before (left) and after capsaicin (right). Tuning vectors are calculated as the mean of all pooled cells in each cluster. Ex1_TACR3/1_ = 57 cells, Ex2_TACR1_ = 75 cells, Ex3_GRPR/NMBR_ = 57 cells, Ex4_CCKBR_ = 21 cells, Ex5_TRHR_ = 65 cells, Ex6_CHRM3_ = 49 cells, and Ex7_OXTR_ = 74 cells from 4 mice.

**Supplemental Fig. 1.**
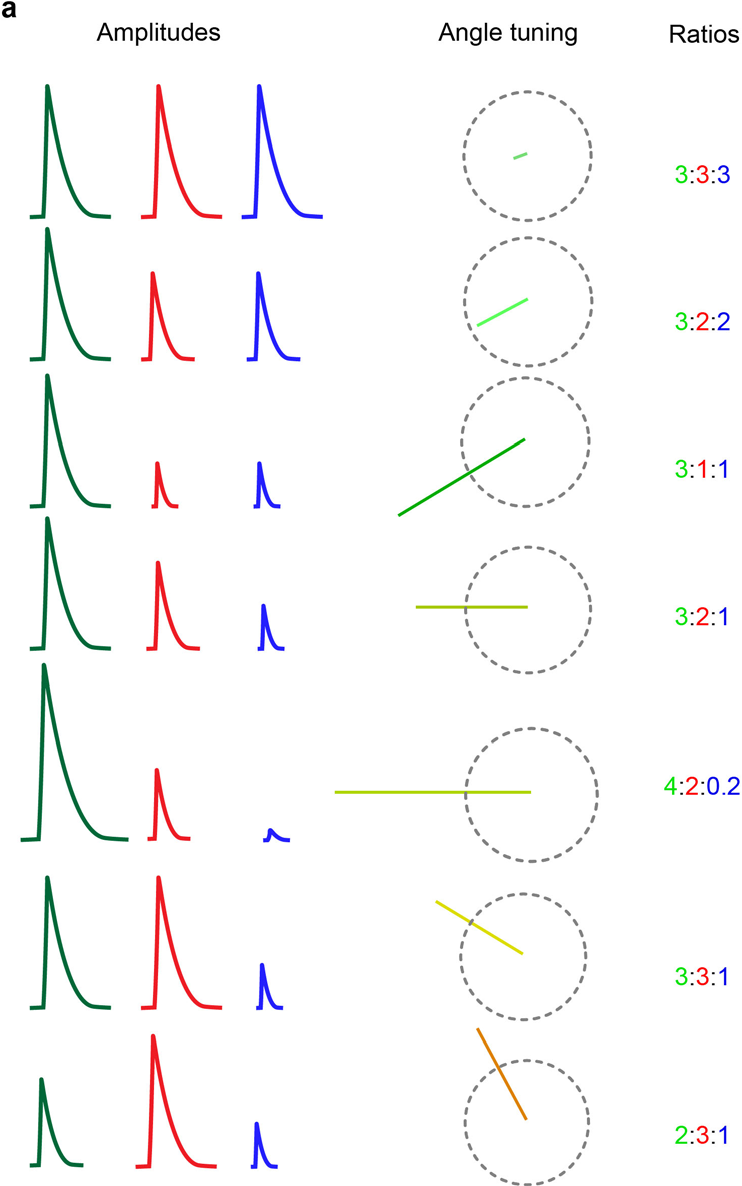
Interaction between amplitude of response and vector tuning angle and magnitude. **a**, Examples of amplitudes of response (left), the ratios of response (right) and the resultant vector tuning for cardinal modalities (mechanical, green; red, heat; blue, cold). Multiple examples are given in order to exemplify how the angle and magnitude represent different tunings. *e.g*. the first 3 examples all have the same mechanical (green) response but as the relative ratios of mechanical to heat/cold are increased, the magnitude of the tuning is increased with maintaining the same angle indicating a stronger preference for mechanical relative to the other modalities permitting an assessment of preference within a polymodal cell.

**Supplementary Fig. 2.**
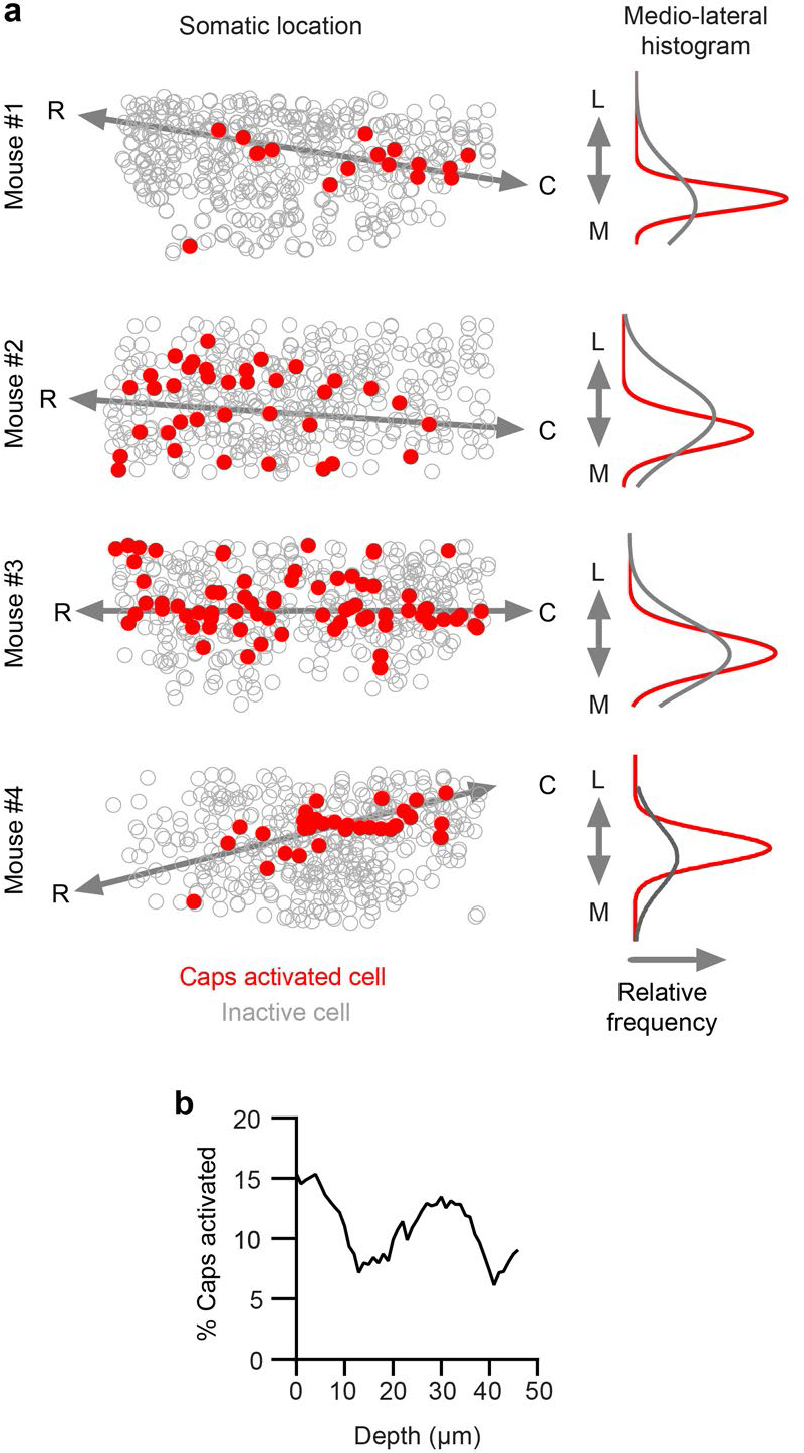
Primary Responders by mediolateral and dorso-ventral distribution. **a**, Left: Somatic location in the SDH of neurons which were either quiescent (grey) or activated (red) after capsaicin injection. Right: The relative frequency of capsaicin activated (red) or all cells (grey). **b**, Percent of cells that were classified as being activated by capsaicin injection (>150% of pre-capsaicin levels based on a 2-minute average of ΔF/F values) across depth.

## References

1. Latremoliere, A. and C.J. Woolf, Central sensitization: a generator of pain hypersensitivity by central neural plasticity. J Pain, 2009. 10(9): p. 895–926.

2. Quesada, C., et al., Human surrogate models of central sensitization: A critical review and practical guide. Eur J Pain, 2021. 25(7): p. 1389–1428.

3. LaMotte, R.H., et al., Neurogenic hyperalgesia: psychophysical studies of underlying mechanisms. J Neurophysiol, 1991. 66(1): p. 190–211.

4. Simone, D.A., et al., Neurogenic hyperalgesia: central neural correlates in responses of spinothalamic tract neurons. J Neurophysiol, 1991. 66(1): p. 228–46.

5. LaMotte, R.H., L.E. Lundberg, and H.E. Torebjork, Pain, hyperalgesia and activity in nociceptive C units in humans after intradermal injection of capsaicin. J Physiol, 1992. 448: p. 749–64.

6. Torebjork, H.E., L.E. Lundberg, and R.H. LaMotte, Central changes in processing of mechanoreceptive input in capsaicin-induced secondary hyperalgesia in humans. J Physiol, 1992. 448: p. 765–80.

7. Baumann, T.K., et al., Neurogenic hyperalgesia: the search for the primary cutaneous afferent fibers that contribute to capsaicin-induced pain and hyperalgesia. J Neurophysiol, 1991. 66(1): p. 212–27.

8. Sluka, K.A., et al., Inhibitors of G-proteins and protein kinases reduce the sensitization to mechanical stimulation and the desensitization to heat of spinothalamic tract neurons induced by intradermal injection of capsaicin in the primate. Exp Brain Res, 1997. 115(1): p. 15–24.

9. Ran, C., M.A. Hoon, and X. Chen, The coding of cutaneous temperature in the spinal cord. Nat Neurosci, 2016. 19(9): p. 1201–9.

10. Chisholm, K.I., et al., Encoding of cutaneous stimuli by lamina I projection neurons. Pain, 2021. 162(9): p. 2405–2417.

11. Todd, A.J., Identifying functional populations among the interneurons in laminae I-III of the spinal dorsal horn. Mol Pain, 2017. 13: p. 1744806917693003.

12. Shortland, P., C.J. Woolf, and M. Fitzgerald, Morphology and somatotopic organization of the central terminals of hindlimb hair follicle afferents in the rat lumbar spinal cord. J Comp Neurol, 1989. 289(3): p. 416–33.

13. Brown, P.B. and J.L. Fuchs, Somatotopic representation of hindlimb skin in cat dorsal horn. J Neurophysiol, 1975. 38(1): p. 1–9.

14. Koerber, H.R. and P.B. Brown, Somatotopic organization of hindlimb cutaneous nerve projections to cat dorsal horn. J Neurophysiol, 1982. 48(2): p. 481–9.

15. Russ, D.E., et al., A harmonized atlas of mouse spinal cord cell types and their spatial organization. Nat Commun, 2021. 12(1): p. 5722.

16. Demsar, J., et al., Orange: Data Mining Toolbox in Python. Journal of Machine Learning Research, 2013. 14: p. 2349–2353.

17. Hachisuka, J., et al., Semi-intact ex vivo approach to investigate spinal somatosensory circuits. Elife, 2016. 5.

18. Micallef, L. and P. Rodgers, eulerAPE: drawing area-proportional 3-Venn diagrams using ellipses. PLoS One, 2014. 9(7): p. e101717.

19. van der Maaten, L. and G. Hinton, Visualizing Data using t-SNE. Journal of Machine Learning Research, 2008. 9: p. 2579–2605.

20. Wang, F., et al., Sensory Afferents Use Different Coding Strategies for Heat and Cold. Cell Rep, 2018. 23(7): p. 2001–2013.

